# Frontotemporal bursting supports human working memory

**DOI:** 10.1101/2025.07.26.666946

**Authors:** Vladimir Omelyusik, Tyler S. Davis, Satish S. Nair, Behrad Noudoost, Patrick D. Hackett, Elliot H. Smith, Shervin Rahimpour, John D. Rolston, Bornali Kundu

## Abstract

Cortical neural activity varies dynamically during memory periods, when relevant information is not present in the environment. But how those dynamics are related to a code defining working memory (WM) performance is not known. Recent data shows brief bursts of activity in the high gamma (70-140 Hz) and beta (12-30 Hz) band within non-human primate lateral prefrontal cortex (PFC) is associated with WM processing. However, WM may be related to activity within a network of frontal executive and posterior sensory areas involved in stimulus perception. Here we tested whether gamma and beta bursting exist in lateral PFC and multisensory lateral temporal areas in humans during visual WM, and whether these areas are coupled via a phase-burst code. We used intracranial macroelectrode recordings from the middle frontal gyrus (MFG), which includes dorsolateral PFC, and from the middle temporal gyrus (MTG), an area important for visual processing. High gamma bursting increased in human left PFC during encoding and delay periods while beta bursting decreased. Interestingly, beta bursting increased in multisensory areas during encoding and remained high during the delay period, more so on the right. These effects varied with WM performance. Finally, we quantify the degree to which delay-period gamma bursting is locked to beta phase within and between regions of this network using a proposed metric termed ‘phase-burst coupling’ (PBC). We find evidence that delay-period gamma bursting in temporal areas is locked to beta phase in PFC. Our findings suggest that WM may use bursting to support memory maintenance until readout.

**Significance Statement:** Working memory is critical for everyday functioning and its disruption results in significant disability in patients with dementia, traumatic brain injury, and stroke. We recorded human intracranial data during a visual working memory task. We found that prefrontal cortex and multisensory temporal brain areas show neural bursting activity during the memory period and that such bursting is related to trial performance. Furthermore, these two areas may be coupled by a phase-burst code. Bursting has been found in non-human primates, rodents, and even crows, suggesting it is a general feature of memory, and perhaps of other aspects of cognition.

## INTRODUCTION

Persistent neural firing has been posited to be the dominant mechanism underlying working memory (WM) processing (1–3). For example, studies in nonhuman primates (NHPs) (4–7) and humans (8) showed that encoding and maintenance of information in memory are associated with sustained activity in single units. However, critical re-analyses of such data have provided evidence for a more dynamic WM mechanism involving transient bursts of neural firing which may be evidence of a population code for mnemonic information itself or its related processing. For example, LFP data from the prefrontal cortex (PFC) of NHPs show sustained elevated gamma band (50-120 Hz) power during the delay period when calculated at the trial-average level; however, at the single-trial level, there are short bursts of increased gamma band and decreased beta band (20-35 Hz) activity (9–12), not sustained delay-period power. Moreover, gamma bursting coincides spatially with single unit neural spiking in relation to memory items, suggesting that bursting codes mnemonic information (13). This was supported by multi-unit recordings in NHPs which show spatial patterns of spiking coinciding with ‘on’ and ‘off’ states in lateral PFC during the delay period, akin to bursting (14). Another study in human middle temporal gyrus showed there are consistent neural firing patterns within a burst, and that those patterns code for item category during a perception task (13). Whether bursting in lateral PFC is relevant in human WM is an open question.

While lateral PFC seems crucial for memory processing (15, 16), WM likely involves other brain regions. The ‘sensory recruitment theory of WM’ posits that memories are maintained through engagement of both PFC as well as the sensory cortices in which the memoranda were originally perceived (17–24) although the underlying mechanism(s) involved have remained elusive. Data from NHPs show that parietal lobe in the context of spatial memory (25) and higher order visual areas (e.g., inferior temporal areas) in the context of object WM (26, 27) show activity related to memory information. Additionally, disrupting delay period activity in sensory areas disrupts memory performance, e.g., see recent transcranial magnetic stimulation study in humans (28). Frontotemporal oscillatory coupling-based mechanisms may contribute to WM processing (3, 29, 30). For example, beta band phase locking between the frontal-eye field area of PFC and the inferotemporal area was related to WM performance in NHPs (31). Here we investigate the role of high gamma and beta band bursts in lateral PFC and in visual/multisensory lateral temporal areas as well as the coupling between and within these areas during object WM in the context of bursts using intracranial recordings from humans.

## RESULTS

We recorded human intracranial data from 7 subjects at the University of Utah (**Table 1**) while they performed an object WM task (**Figure 1a**). These subjects were patients with epilepsy who had depth electrodes implanted for clinical seizure monitoring and hospitalized during that time. All participants achieved satisfactory performance: 3 of 3 items correct: 56% ± 19%, 2 of 3 items correct: 31% ± 11%, 1 of 3 items correct: 11% ± 8%, 0 of 3 items correct: 2% ± 1%). Trials with 3 of 3 items correct are referred to here as ‘correct’ trials and trials with 0 or 1 item correct as ‘incorrect’ trials (**Figure 1b**).

**Figure 1.**
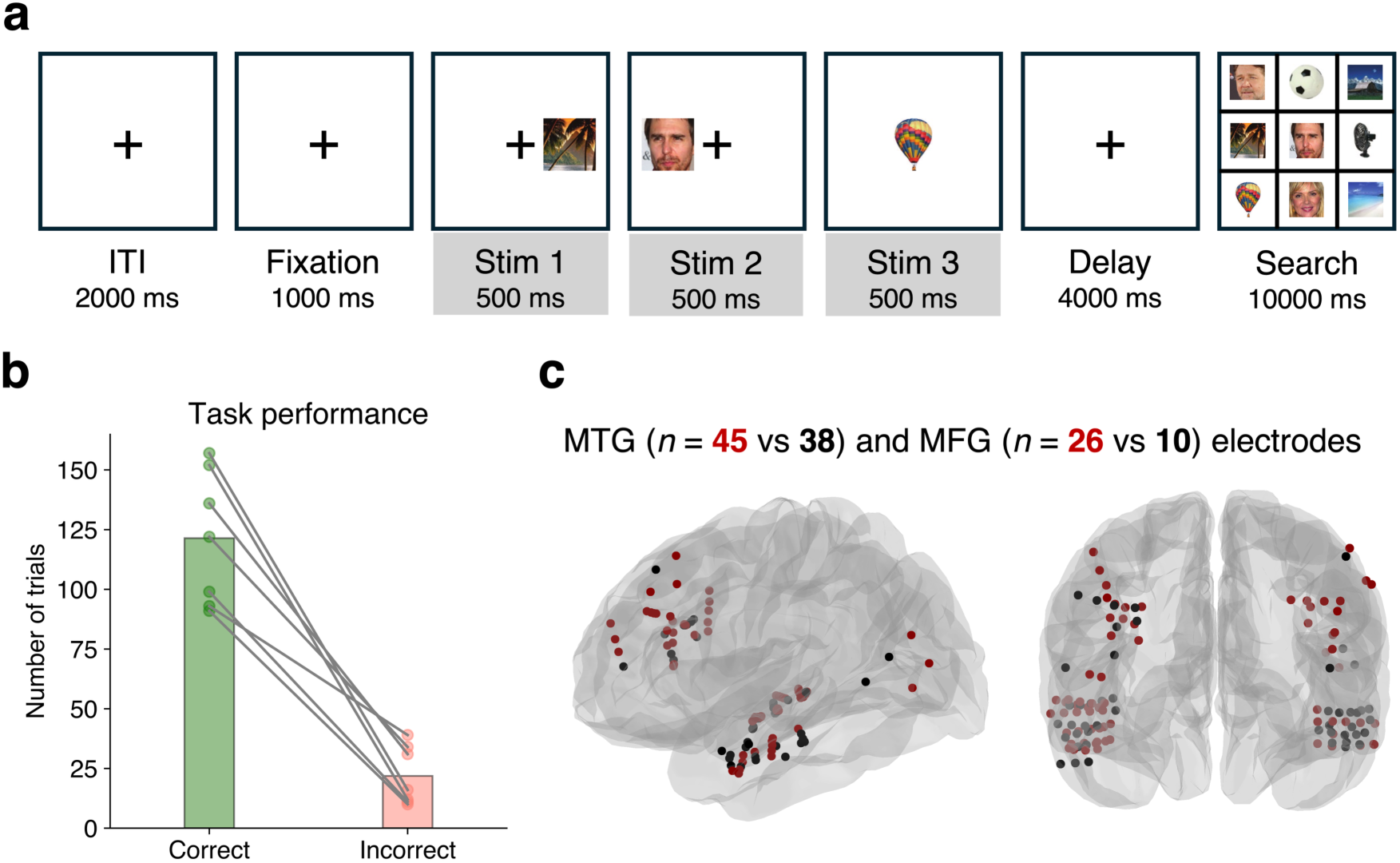
Task and behavioral results. **a.** Object working memory task. A fixation period (1000 ms) was followed by 3 consecutive visual stimuli (500 ms each) from one of 3 categories (faces, objects, scenes). Items were presented in the center, left, or right position. After a delay period (4000 ms), the subjects searched for the target stimuli in a grid of 9 items (up to 10 sec), followed by an inter-trial-interval (ITI, 2 sec). **b.** Task performance across 7 subjects from the Utah cohort. Trials which had 3 of 3 items chosen correctly were labeled as ‘correct’ and trials which had 0 or 1 items chosen correctly were labeled as ‘incorrect’. Circles correspond to individual subjects’ performance; bars represent the average. **c.** Spatial distribution of broadband-modulated (red) and non-modulated (black) channels within ROIs, across subjects.

**Table 1.**
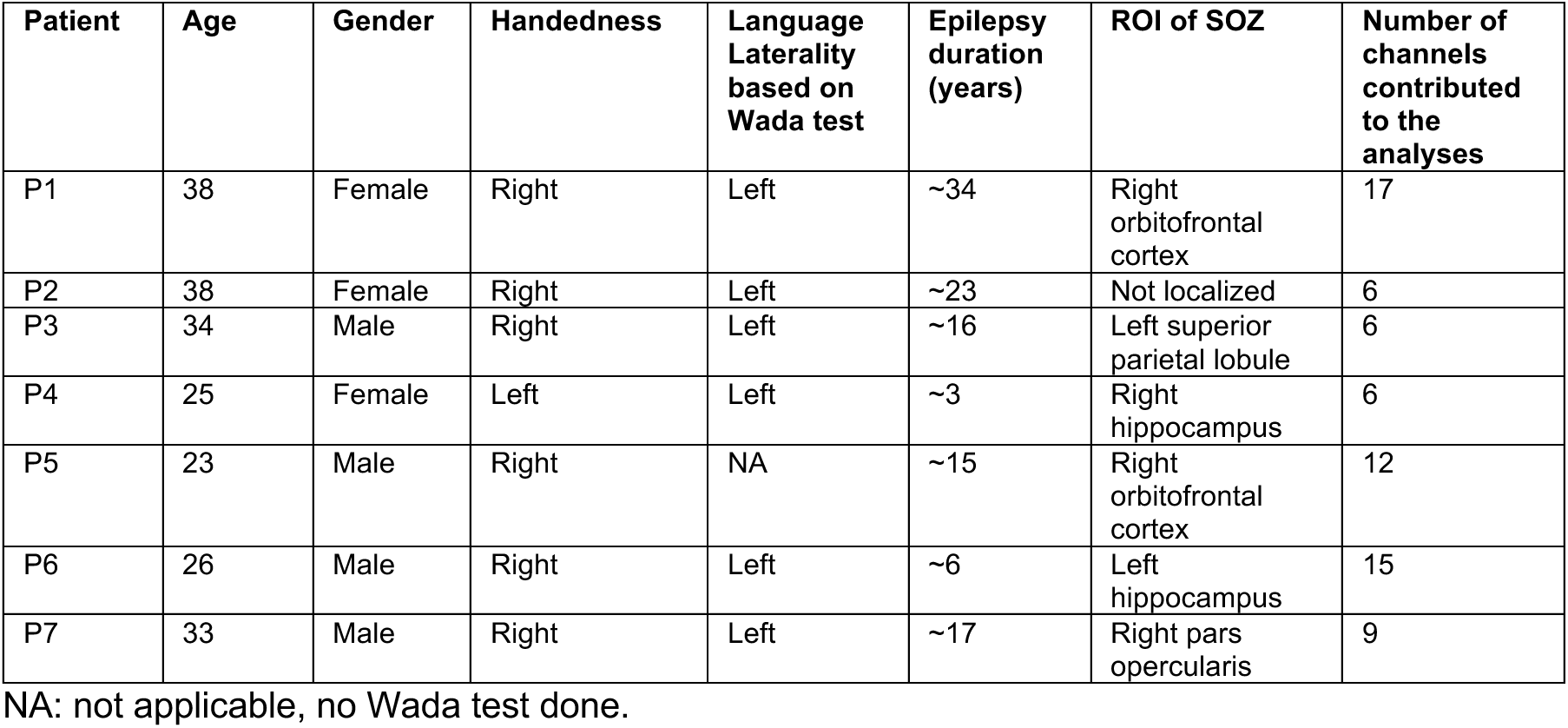
Patient demographics and relevant characteristics (University of Utah cohort)

Four regions of interest (ROIs) were analyzed: the left and right middle frontal gyri (LMFG/RMFG, collectively referred to as PFC) and the left and right middle temporal gyri (LMTG/RMTG) which are multisensory areas (32, 33). These frontal and temporal areas were chosen because they were relevant to WM and adequately sampled in this experimental setup. In each ROI, channels with significant broadband (30–140 Hz) activity during the encoding period of the first stimulus in the trial were identified. This activity is a feature related to stimulus perception and memory encoding (34). Further analyses were restricted to these channels (**Figure 1c, Figure S1, Table 2**). Overall, 84% of channels in PFC and 83% of channels in MTG showed significant stimulus-evoked activity.

**Table 2.**
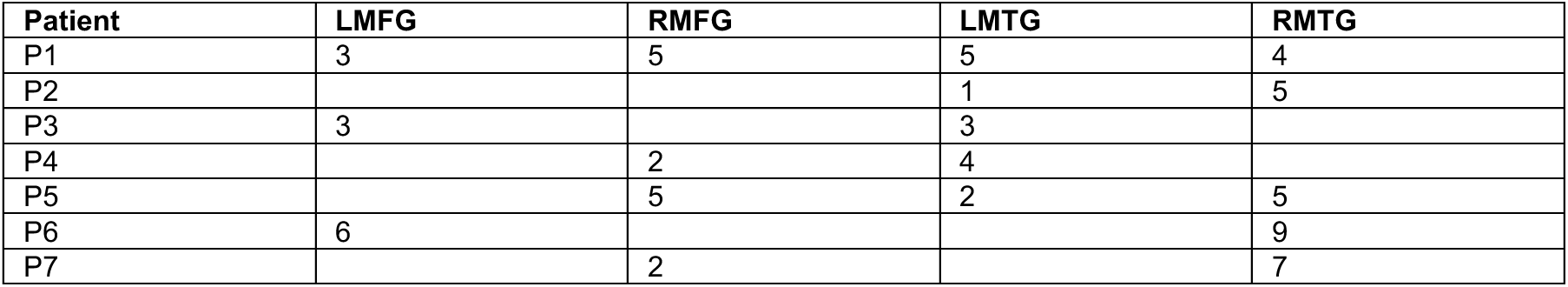
Location of channels (University of Utah cohort)

### Humans show bursts of power in PFC during WM

Sustained high gamma (70–140 Hz) and beta (12–30 Hz) power changes were observed in trial averages during encoding and delay periods, relative to fixation, in left dorsolateral PFC. Trial-averaged beta power in LMFG was suppressed during encoding and delay periods, relative to fixation, and high gamma power was elevated relative to fixation (**Figure 2a**; *p* < 0.05, cluster test). Single trials, however, revealed transient bursts of power rather than sustained increases or decreases in power during the encoding and delay periods (**Figure 2b, Figure S2a**). We defined ‘bursts’ as periods during which power exceeded 2 standard deviations (SD) above baseline for at least 3 oscillatory cycles (9). Beta and high gamma bursts were detected in all ROIs during all time periods (**Figure S3**). Mean delay-period trial-by-trial burst rates were 1.7 ± 0.98 bursts per second for high gamma bursts and 0.04 ± 0.02 bursts per second for beta bursts (**Methods**).

**Figure 2.**
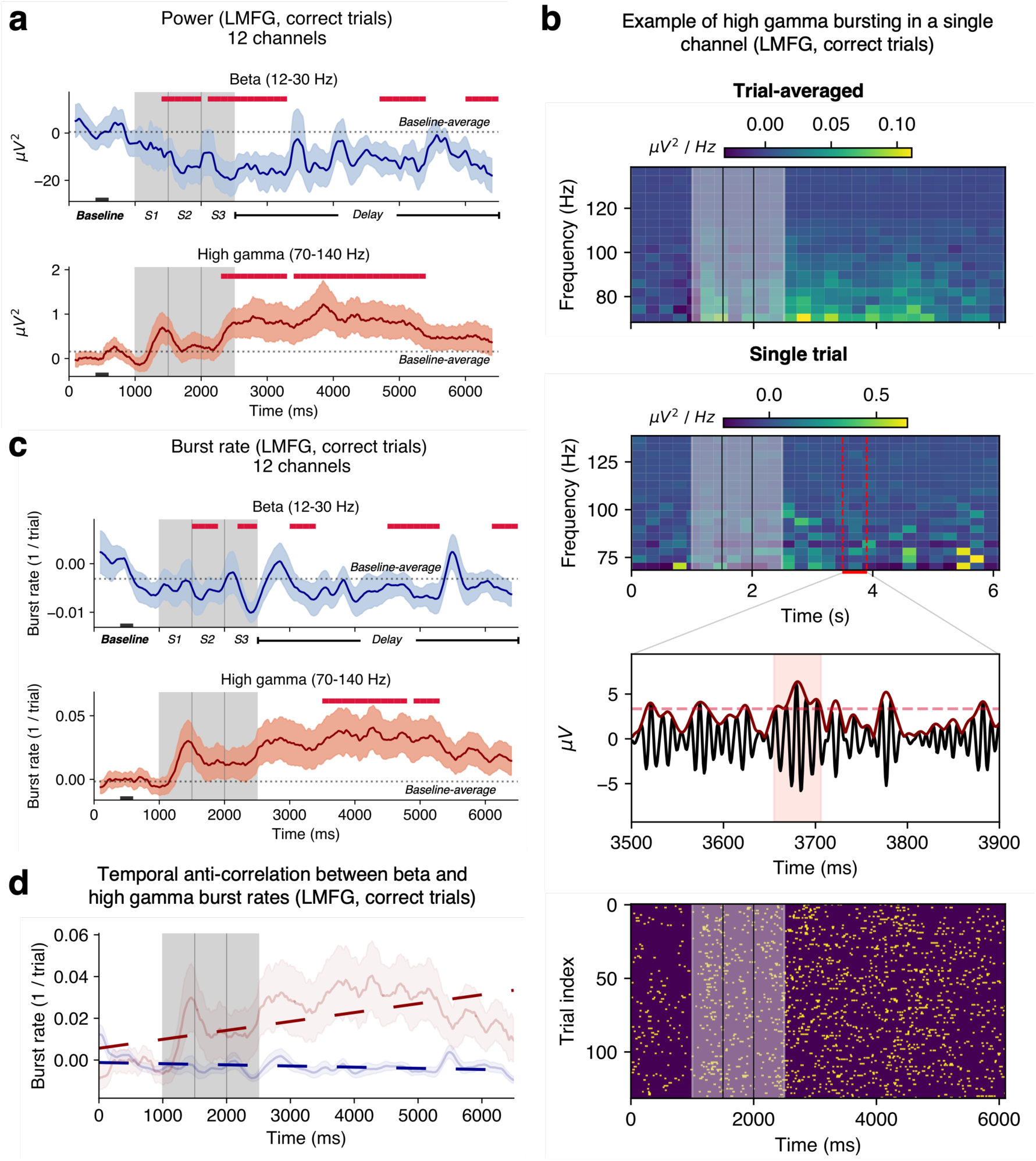
Human WM is associated with bursting in LMFG (correct trials, 12 channels, 3 subjects) **a.** Beta and high gamma power averaged over channels. **b.** Top two panels: examples of trial-averaged and single trial spectrograms in the high gamma frequency band (70-140 Hz) from one representative channel. Trial-averaged power is sustained over encoding and delay periods, but the single-trial spectrogram shows short bursts of elevated power. **Inset**: burst detection within trial (raw LFP in black, analytical amplitude in dark red; dashed line is the threshold for burst detection; the shaded red region corresponds to a detected high gamma burst). **Bottom panel**: burst raster for that channel, encoding period in grey. **c.** 1/trial beta and high gamma burst rates. **d.** Anticorrelation between high gamma and beta burst rates over the trials. In panels **a, c, and d**, shaded regions correspond to ±1 SEM. Crimson lines indicate significant differences from the mean rate during 400-600 ms of fixation denoted with black rectangle at the bottom (*p* < 0.05, cluster test; 200 ms running window, 50% overlap). A 200 ms smoothing window was used for visualization but not for testing.

While burst rate can be calculated trial-by-trial as above, it can also be assessed through a trial-aggregate ‘1/trial’ burst rate (**Methods**) which is the proportion of trials with a burst at that time point in the trial scheme (9). Throughout, this 1/trial burst rate is referred to as ‘bursting’ or ‘burst rate’ and we specify instances of the ‘trial-by-trial burst rate’ explicitly. In LMFG, similar to trial-averaged power measures, there was decreased beta bursting during the encoding and delay periods, and increased high gamma bursting during the delay period relative to fixation (**Figure 2c**). Trends in RMFG differed from those in LMFG. Trial-averaged beta power decreased relative to fixation only during encoding and there was no change in high gamma power compared to baseline (**Figure S4a**). While beta and high gamma bursts were detected in this area (**Figure S2b, Figure S3, Figure S4b**), their rates did not change during the trial (**Figure S4c**).

In LMFG, beta and gamma burst rates were negatively correlated over the full trial duration (average *r* = −0.08 with *p* < 0.05 for 9/12 channels, **Figure 2d**). However, over short time windows, individual channels within an ROI showed both significant positive and negative correlations (150 ms duration windows, **Figure S5a**). This effect was not patient-specific (*p*s < 0.05, 1-way ANOVA at each time point) and may suggest there exists a dynamic population code that changes over short-time scales, and which manifests as fluctuations in bursting activity (2, 9).

### Bursting also occurs in the human lateral temporal lobe during WM

Trial-averaged power dynamics differed in the multisensory areas compared to that in the PFC. High gamma power was suppressed in RMTG only during encoding with no change in beta power, relative to fixation (**Figure 3a**; *p* < 0.05, cluster test). High gamma and beta bursts were detected in the right and left temporal regions (**Figure 3c-d, Figure S6b**). In contrast to trends in LMFG, delay-period beta burst rate in RMTG was *elevated* during encoding and delay periods relative to fixation (**Figure 3c**, *p* < 0.05, cluster test). Mean delay-period trial-by-trial burst rates were 0.72 ± 0.23 bursts per second for high gamma bursts and 0.19 ± 0.1 bursts per second for beta bursts (**Figure S3**). In LMTG, there was also an increase in beta band burst rate at the end of the delay period relative to fixation (**Figure S6c**). Mirroring the finding in LMFG, RMTG high gamma and beta burst rates were significantly anti-correlated over the trial scheme (**Figure 3d**; average *r* = −0.04 with *p* < 0.05 for 24/30 channels) but not when considering smaller 150 ms time windows (**Figure S5c**).

**Figure 3.**
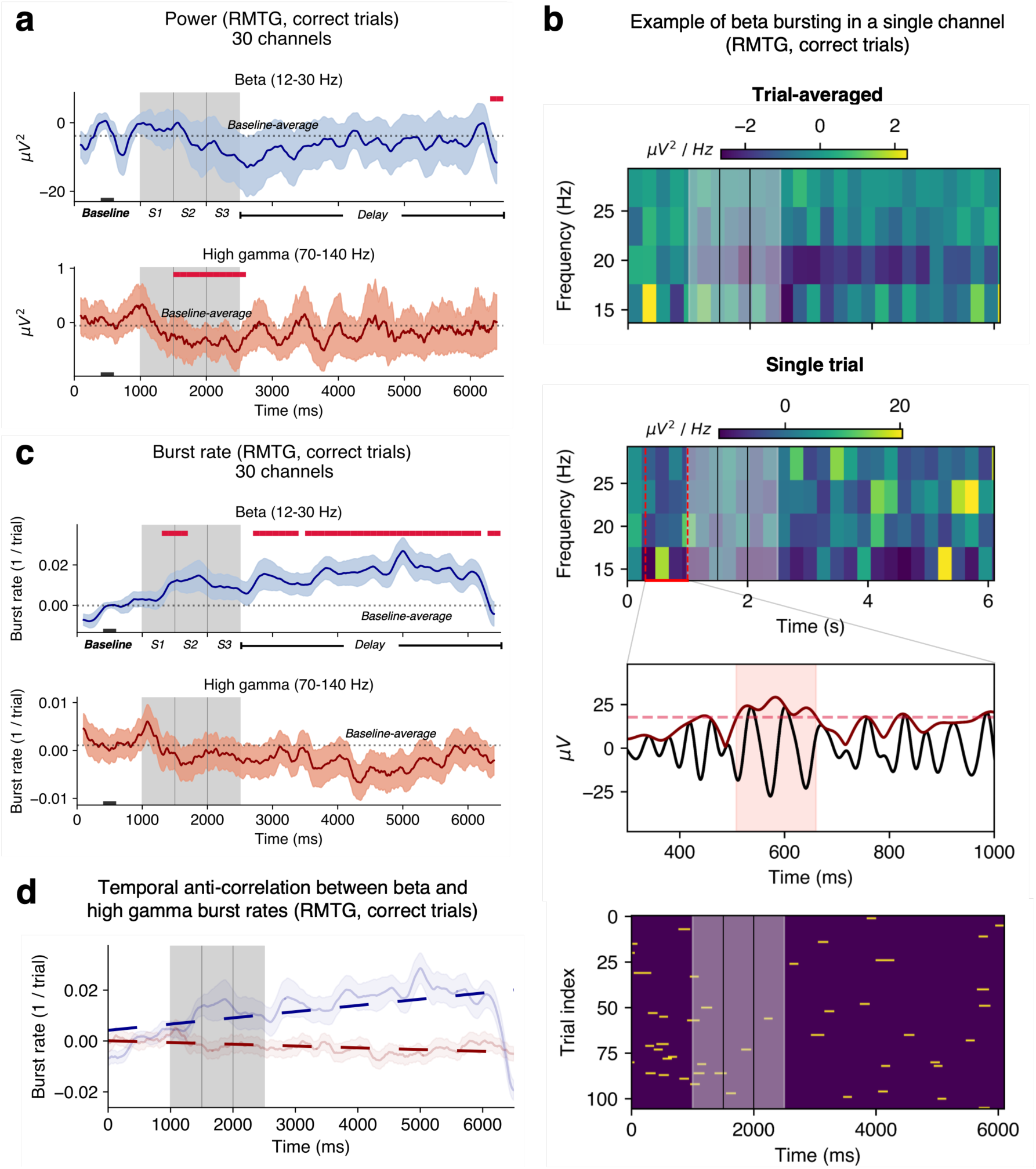
WM-related bursting is seen in RMTG (correct trials, 30 channels, 5 subjects) **a.** Beta and high gamma power averaged over channels. **b.** Top two panels: an example of trial-averaged and single trial spectrograms in the beta frequency band (12-30 Hz) from one representative channel. **Inset**: an example of burst detection within that trial (raw LFP in black, analytical amplitude in dark red; dashed line depicts the threshold used for burst detection; the shaded red region corresponds to a detected beta burst). **Bottom panel**: burst raster for that channel, encoding period in grey. **c.** 1/trial beta and high gamma burst rates. **d.** Anticorrelation between high gamma and beta burst rates over the trials. In panels **a, c, and d**, shaded regions correspond to ±1 SEM. Crimson lines indicate significant differences from the mean rate during 400-600 ms of fixation (black rectangle at the bottom; *p* < 0.05, cluster test; 200 ms running window, 50% overlap). A 200 ms smoothing window was used for visualization but not for testing.

### Burst rates in left PFC and multisensory temporal areas vary with task performance

In the left dorsolateral PFC, there was a significant increase in high gamma bursting during the delay period and a decrease in beta bursting during the encoding periods for correct compared to incorrect trials (**Figure 4a***, p* < 0.05, cluster test). These gamma and beta results were qualitatively reflected in the period-averaged burst rate data (**Figure 4b**; 2-way GLME, no significant effects of performance, task period or task x period interaction found for either burst type; post-hoc pairwise tests, Tukey correction) and in the trial-by-trial burst rates (**Figure S3**) although not all comparisons were significant. In RMTG, there was a significant increase in beta bursting for correct trials compared to incorrect trials mainly during the delay period (**Figure 4a**; *p* < 0.05, cluster test). Period-averaged burst rates reflected the beta bursting effect qualitatively (**Figure 4b**; 2-way GLME, high gamma bursts: no significant effects found, beta bursts: no significant interaction between period x performance, no main effect of performance, main effect of period, *F* = 3.47, *DF1* = 2, *DF2* = 174, *p* = 0.03; post hoc pairwise tests, Tukey correction) although not all comparisons were significant. Similar results were found using the raw beta burst rates in RMTG (**Figure S7a**). These results show that bursting in left dorsolateral PFC (LMFG) and right multisensory area (RMTG) may influence performance on this WM task.

**Figure 4.**
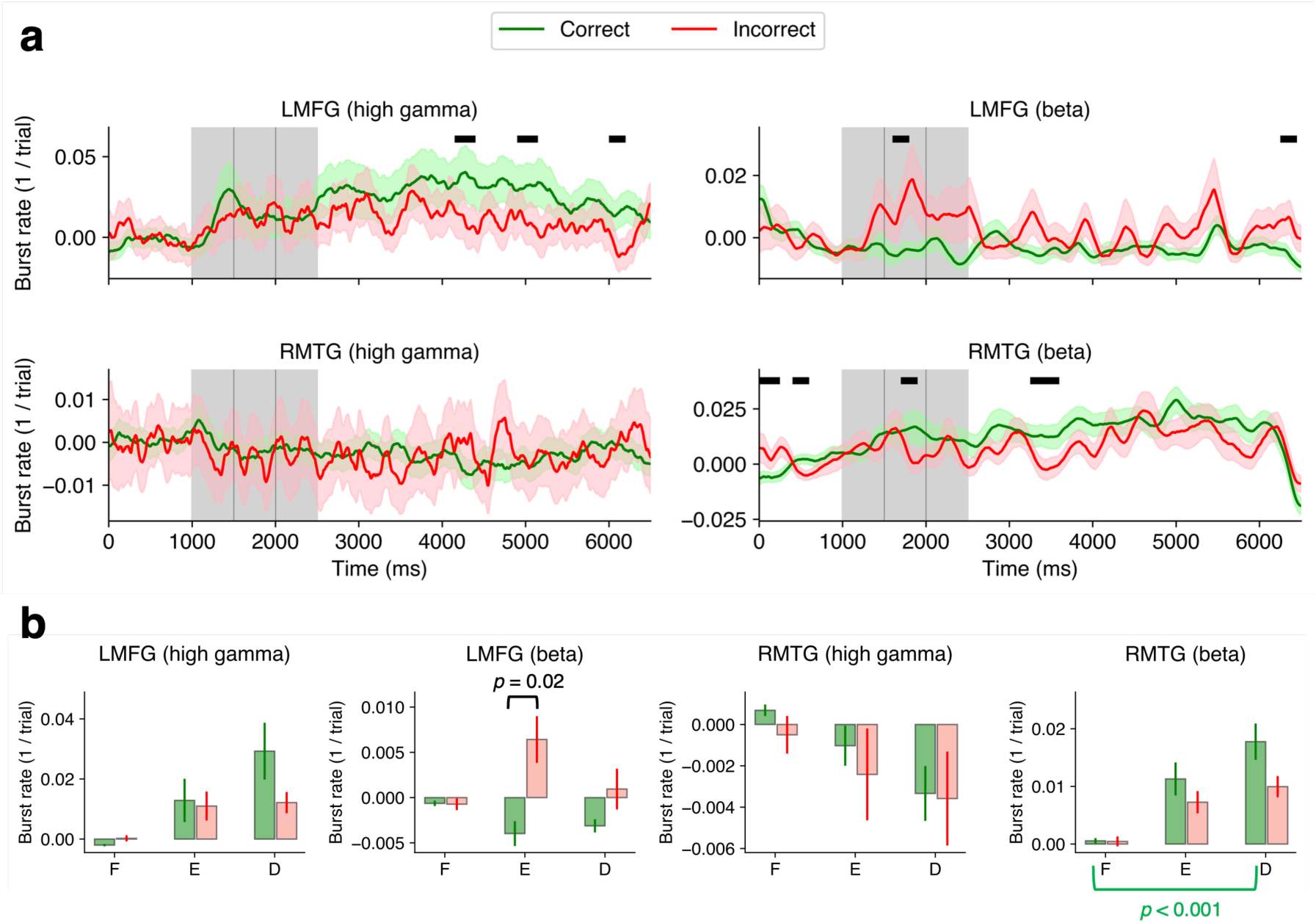
Burst rates in LMFG and RMTG differentiate WM performance. **a.** Baseline-subtracted burst rates for correct (green) and incorrect (red) trials. Shaded regions represent ±1 SEM. Black lines correspond to periods of significant difference between correct and incorrect averages over channels (*p* < 0.05, cluster test; 200 ms running window, 75% overlap). The baseline period was 200-800 ms from the start of the trial. A 200 ms smoothing window was used for visualization but not for testing. **b.** Bar plots of burst rates during 3 periods (F-fixation, E-encoding, D-delay) and 2 performance conditions (correct/incorrect trials). 2-way GLME, **LMFG**: no significant effects of performance, period or their interaction found for either burst type; **RMTG**: high gamma bursts: no significant effects found, – beta bursts: no significant interaction between period x performance, no main effect of performance, main effect of period, *F* = 3.47, *DF1* = 2, *DF2* = 174, *p* = 0.03; post-hoc pairwise tests, Tukey correction). Error bars correspond to ±1 SEM.

### Bursting in PFC is coupled with bursting in multisensory areas

Interestingly, high gamma and beta burst rates changed in parallel over the trial between PFC and multisensory temporal areas (**Figure S8**). We hypothesized that this effect might be related to coupling of beta and high gamma rhythms between frontal and temporal regions. Evidence from NHP and human data shows that PFC may exert beta-band mediated ‘top-down’ executive control onto other brain regions during various cognitive tasks (35–38). On the other hand, sensory areas may be recruited for memory representation and propagate information ‘bottom-up’ in an automated fashion, through the ventral visual stream (39). We investigated possible network linkages via several traditional metrics and a proposed metric, PBC, that captures such coupling.

#### Directional phase lag index (dPLI)

This measure (40) can quantify whether the frontotemporal phase differences occur in a consistent direction over trials, i.e., if one region follows the other consistently over the trial scheme. Here, dPLI is 0.5 if PFC consistently leads multisensory areas, 0 if there was no consistent leader, and −0.5 if multisensory areas lead PFC. We found that LMFG led multisensory RMTG (**Figure 5a**). There was a significant dPLI elevation in the beta band during the delay period in correct trials. No significant leader was found during incorrect trials for most of the delay period until RMTG led in the last 1.5 seconds prior to search.

**Figure 5.**
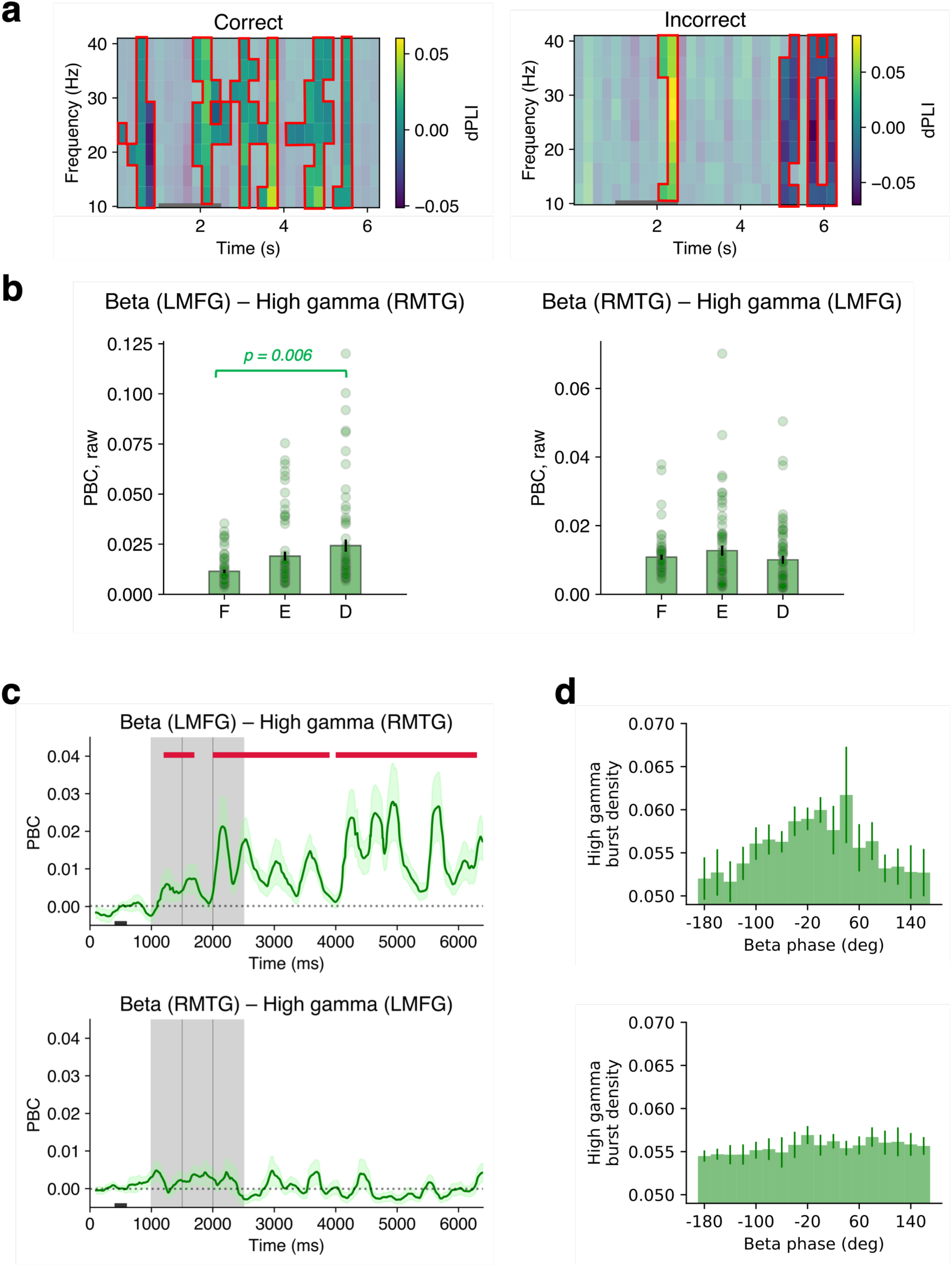
LMFG and RMTG are linked through phase-burst coupling (PBC) (66 pairs between LMFG and RMTG, 2 subjects) **a.** Beta-beta dPLI (directed phase lag index) between channels in LMFG and RMTG. Scale bar: yellow indicates LMFG leading in phase, blue indicates RMTG leading in phase, non-significant cells are greyed out (*p* < 0.05, cluster test). **b.** Raw beta-high gamma PBC between LMFG and RMTG without baseline-subtraction. Delay-period PBC in the LMFG-RMTG pair was significantly different from fixation (pairwise post-hoc tests, Tukey correction). Error bars correspond to 1 SEM. **c**. Baseline-subtracted beta-high gamma PBC between (**top**) LMFG and RMTG; (**bottom**) RMTG and LMFG. Crimson lines indicate significant differences from the mean PBC from 400-600 ms of fixation indicated by black rectangle at the bottom (*p* < 0.05, cluster test; 200 ms running window, 50% overlap). A 200 ms smoothing window was used for visualization but not for testing. **d. (Top)** Distribution of high gamma bursts during the delay period of RMTG over the beta phase of LMFG. **(Bottom)** Distribution of high gamma bursts during the delay period of LMFG over the beta phase of RMTG. Green error bars correspond to ±1 SEM.

#### Phase-burst coupling

We then tested whether high gamma bursts from one ROI occur at specific phases of the underlying beta rhythm of another ROI. Such consistency in burst positioning may be a mechanism to informationally link areas showing a signal phase lag. We defined a synthetic measure termed phase-burst coupling (PBC) value, adapted from the commonly used modulation index to include the time dimension, see (41–43). The PBC value compares the entropy of the test high gamma burst distribution over all beta phase bins to that of a uniform distribution in a moving window. This measure reflects the degree to which bursts cluster at particular phases of the ongoing underlying oscillation. The measure ranges between 0 and 1 with higher values indicating more clustering or ‘modulation’. Note that this measure is sensitive to the number of cycles considered (42), see **SI Methods and Results.** For this reason, we report cycle-balanced results throughout for various comparison conditions. We calculated PBC of beta phase and high gamma bursts between LMFG and RMTG channel pairs, constructing phase-burst distributions over a 150-ms (∼ 3 beta cycles) window and over trials for each time point (**Figure 5b).** Throughout, we refer to the oscillation used for the phase variable first, followed by the oscillation used for the burst variable second, e.g. ‘beta – gamma PBC’ refers to beta oscillatory phase in ROI-1 coupled to gamma oscillatory bursts in ROI-2. Thus, the measure has built in directionality. It can also be calculated within an ROI.

Beta–high gamma PBC between left PFC and RMTG was significantly elevated during the encoding and delay periods, compared to fixation, in correct trials (i.e. ‘top-down’ effect; **Figure 5c**; *p* < 0.05, cluster test, data from 2 patients). Interestingly, beta–high gamma PBC between RMTG-LMFG, which is in the opposite anatomical direction, remained at baseline (i.e. ‘bottom-up’ effect). These PBC results, although limited by the amount of data at hand, suggest consistent patterns of high gamma burst positioning on the phase of beta oscillations during the delay period. Considering frontotemporal channel-pairs that showed a significant degree of non-uniformity (on average, 13% of channel-pairs, *p* < 0.05, Rayleigh test, Bonferroni-corrected; **Figure 5d**), we found that the gamma bursts in RTMG were at the peaks of beta phase in LMFG during correct trials. Namely, the modes of the distributions varied between directional ROI pairs (top 2 modes, degrees: beta (LMFG) – high gamma (RMTG): 40, 0; beta (RMTG) – high gamma (LMFG): −20, 80. Finally, frontotemporal beta-high gamma PBC between left PFC and RMTG was elevated for correct compared to incorrect trials during the delay period (**Figure S9b**; *p* < 0.05, cluster test). Thus, gamma bursts occurred at different beta phases between frontotemporal ROIs, suggesting that PBC may enable information communication over long distances at the scale measured by macroelectrodes.

We also explored whether the PBC metric captured coupling relationships between other ROI pairs and found that delay period beta–high gamma PBC between LMFG-LMTG and within LMTG was briefly elevated relative to fixation during correct trials (**Figure S10a,b**; *p* < 0.05, cluster test). It may be the case that not all pairs of channels measured are relevant to WM, so we tested WM performance effects in channels with elevated PBC (top 20%) during correct trials. While the data are also limited, beta–high gamma PBC between RMTG-LMFG, and within both RMTG and LMTG were elevated for correct compared to incorrect trials (**Figure S11**; *p* < 0.05, cluster tests for all comparisons).

#### Mutual Information

We also calculated mutual information trial-by-trial between high gamma bursts of LMFG and RMTG channels (see **SI Methods and Results**, (44)). There was no effect of task period or WM performance on mutual information (**Figure S12**; 2-way GLME, all *p*s>0.05), suggesting there is no difference in the amount of information present in gamma burst timing between frontal and temporal ROIs during the delay period, compared to fixation, across all available channels.

#### Phase-phase locking (PPL)

We computed the trial-by-trial PPL between PFC and MTG and the phase locking value (PLV) measure within ROIs (see **SI Methods and Results**, (31, 45, 46). There was a significant decrease in frontotemporal beta PPL between LMFG and RMTG during the second half of the delay period for correct trials (**Figure S13a**; *p* < 0.05, cluster test) and an increase in delay period beta PPL for incorrect vs correct trials (**Figure S13b**). Notably, there was a significant increase in beta PLV within RMTG during both encoding and delay period for correct trials (**Figure S13a**).

### WM task variation tested in a separate cohort of patients

We recorded data from 6 additional patients at the University of Missouri with a similar experimental setup and protocol as the Utah cohort except that subjects were not required to maintain fixation during encoding (**Table S1, Table S2, Figure S14a-b**). They could foveate to the location of the three stimuli presented left, right, or center, during encoding, but then held fixation during the delay period (based on eye tracker data). Note that this likely changes the nature of the top-down control required by dorsolateral PFC, which has a known role in spatial WM and spatial attention (47). Indeed, much of the data from WM tasks in NHP involve object perception in extrafoveal locations during central fixation (48). Patients performed better on this task compared to the Utah cohort (3 of 3 items correct: 76.5% ± 15.2%, 2 of 3 items correct: 20.2% ± 12.9%, 1 of 3 items correct: 3% ± 2.7%, 0 of 3 items correct: 0.3% ± 0.5%; *p* < 0.001, χ^2^ contingency test), which may be related to this difference in task instruction and execution. Due to low numbers of incorrect trials, we limited analysis to correct trials.

Delay-period high gamma burst rate in LMFG (**Figure S12c, left)** and beta burst rate in RMTG were not significantly different from fixation (**Figure S14c, right**, *p* < 0.05, cluster test, correct trials only). However, RMTG beta burst rate showed qualitatively similar trends and amplitudes between cohorts (mean beta burst rate in RMTG was 0.04 in the Utah cohort and 0.04 in the Missouri cohort). Also as in the Utah cohort, high gamma burst rate in LMFG was significantly anticorrelated with the beta burst rate over the task schema (**Figure S15c**; average *r* = −0.04 with *ps* < 0.05 for 9/11 channels), while the relationship was opposite in RMTG (average *r* = −0.008 with *ps* < 0.05 for 8/14 channels). Baseline-subtracted beta-gamma PBC over LMFG and RMTG pairs was not significantly different from fixation (**Figure S14e**). Thus, the influence of dorsolateral PFC onto multisensory areas was not as prominent in this task variation.

## DISCUSSION

WM-related bursting has recently gained attention after re-analyses of classic WM task data showed that PFC neurons more commonly exhibit sparse or dynamic firing trial-by-trial as opposed to persistent firing at a stable rate (1, 2). Gamma LFP derived from micro-electrodes showed bursts of power that spatially and temporally overlap with informative spiking during the delay period in PFC of NHPs (9, 10). At the cortical layer level, microelectrode recordings from NHPs show gamma power is elevated in superficial layers of PFC during attention and memory tasks while alpha or beta power is elevated in deeper layers (49, 50). Furthermore, WM may involve coordinated activity of a network of PFC and sensory and multisensory areas (3, 22–24, 29). Here we investigated whether bursting is related to memory maintenance within a frontotemporal network commonly implicated in WM using macroelectrode recordings which capture large-scale population dynamics within and across brain regions.

Using human intracranial data, we showed that bursting in PFC and in multisensory temporal areas supports visual WM. Specifically, 1) we found increased high gamma burst rate and a concomitant decreased beta burst rate in the left PFC during the delay period, similar to that in NHPs (9, 10). 2) WM-related bursting also existed in multisensory temporal areas with increased beta bursting during the delay period in RMTG. 3) High gamma bursting in PFC and beta bursting in temporal lobe varied with WM performance. Finally, limited data showed that 4) beta-high gamma PBC between frontal and temporal areas as well as within temporal areas varies with WM performance. Together these results indicate that multi-region bursting is related to memory maintenance and performance and that PBC may be a mechanism that supports communication in this network at a macroscopic population level.

### Bursting in PFC and multisensory areas during human WM

Both beta and high gamma bursting in the human PFC predicted WM performance mirroring results from a match/non-match task in NHPs (9). In humans, while the 30-60 Hz power has been shown to increase with memory load during the encoding period, measured with intracranial recordings (51); to our knowledge, gamma bursts have not been reported in human dorsolateral PFC during the WM delay period. We found that in LMFG, the high gamma burst rate was higher for correct trials and scaled with the number of correct responses (**Figure S16**) implying gamma bursting reflects some aspect of memory fidelity.

We also found that the beta burst rate was lower for correct versus incorrect trials in the left PFC (**Figure 4**). Beta activity may reflect functional inhibition such that at an operational level, it disrupts presumably “unneeded” computations (52). In movement planning and execution, several studies have demonstrated an inhibitory role of beta oscillations (53–55). Parkinson’s disease is associated with excessive beta bursting during motor movements, causing bradykinesia (56) and deep brain stimulation is targeted to those areas of the subthalamic nucleus with high beta power. Recording from dorsolateral PFC in Parkinson’s disease patients, another study reported decreased beta power during the encoding period of a WM task (57). In our data, there was a significant decrease in beta power in the bilateral middle frontal gyri during encoding. However, right and left PFC showed different bursting behavior during this WM task. This may indicate different roles for the right and left PFC regions. Right PFC has been implicated in reasoning (58) and interference control, which was not used in this task (59, 60). Left PFC may be more important for maintaining information during the delay period in our task that has no interference or load manipulation components.

We found evidence that WM-related bursting is not unique to the PFC but also exists in the multisensory area middle temporal gyrus, which is involved in both auditory and visual tasks (33). It is a uniquely human area that is presumably an evolutionary extension of the superior temporal sulcus, which is also found in NHPs. RMTG beta burst rate increased during both the encoding and delay periods (**Figure 3c**). Interestingly, this increase in burst rate occurred despite RMTG power remaining at baseline levels during the trial schema (**Fig. 3c** vs **3a**) suggesting that consideration of burst dynamics over general power is relevant to WM function. Beta burst rate in RMTG was higher in correct compared to incorrect trials (**Figure 4**).

Bursts have been recorded in humans in other brain areas during WM. Delay period alpha (8-14 Hz) and beta bursts were recorded in parietal and occipital lobes during human WM, using magnetoencephalography (61). Using human intracranial data during a long-term memory task, the number of high gamma range bursts in the ventral visual stream varied with performance during the encoding period (62). However, this relationship was not explored during the period when the stimulus was not on screen as in our experiment. Peri-hippocampal ripple (70-180 Hz) power increased in the fusiform gyrus in humans just prior to the free recall of memory items in a long-term memory task and preceded hippocampal sharp wave ripple by 18 ms (34). Additionally, frontal and hippocampal ripples discriminated between events and event boundaries during movie watching (63).

Beta and high gamma burst rates in RMTG were anticorrelated during the memory trial, and the trends were opposite to that in bilateral PFC (**Figure S8**). Beta burst rate also increased in LMTG at the end of the delay period in correct trials (**Figure S6c**), and beta and high gamma burst rates exhibited a similar anticorrelation effect as in RMTG. This bilateral MTG anticorrelation effect was reproduced in the Missouri cohort (**Figure S15c**). Increased beta bursting in RMTG during the delay period may indicate suppression of task irrelevant neural ensembles locally. We also found increased beta band PPL within RMTG during the delay period (**Figure S13**). Overall, these results are in line with the sensory working memory hypothesis which posits participation of multisensory areas in mnemonic coding of visual stimuli (22). Note that we did find elevated high gamma burst rate in LMTG for incorrect trials primarily during encoding and delay periods (**Figure S7c**), which may reflect encoding errors. Further work is needed to tease apart the role of bursting in these lateralized temporal areas.

### Burst-mediated coupling as a mechanism in WM

Anticorrelation between the dynamics of high gamma and beta burst rates in PFC and multisensory temporal areas indicates potential frontotemporal coupling supporting a WM network (11, 29). Although our data were limited by the electrode combinations subjects had, we found evidence that LMFG leads RMTG through signaling in the beta band during the delay period for correct trials based on a directional phase locking measure, dPLI. Temporal areas led at the end of incorrect trials, perhaps “ramping up influence” (the ramp up of dPLI at the end of the incorrect trial, **Figure 5a**) to retrieve memories and perform the end search, a general feature which has been noted in other studies (34). Simultaneous recordings from PFC and parietal cortex in NHPs show object identity-specific beta coherence during the delay period and that PFC leads the parietal cortex during the delay period (64).

To explore such coupling further, we developed a phase-burst coupling metric, PBC, to characterize the organization of LFP burst timing relative to the phase of ongoing oscillations between two areas (41, 65). Note that while this analysis was performed ‘over-trials,’ it captures the unique relationship between the underlying oscillatory phase and burst timing that may be more consistent over all trials of the task than, for example, the PPL metric. There was increased coupling between high gamma bursts in temporal areas and beta phase in PFC for correct trials, and bursts were at the peaks of the beta phase (**Figure 5**). Furthermore, delay-period frontotemporal PBC was elevated for correct compared to incorrect trials (**Figure S9**). Overall, PFC seems to provide some level of directional and perhaps top-down control during this task that is specific to memory maintenance, and memory performance. Note that PBC does not necessitate causality. That would have to be tested by systematically changing aspects of the ongoing phase or bursts. Such techniques might include noninvasive and invasive stimulation that have been applied to modulate memory performance (66–69).

Coupling has been measured in various settings recently using invasive and non-invasive field potential recordings along with neuronal spiking data in other frontal and temporal brain areas during WM tasks, but without consideration of bursting. In humans, Daume et al. (2023) used intracranial microelectrode recordings to show coupling between the theta rhythm in ventromedial PFC and hippocampal spikes during the delay period of a similar WM task (70). Using magnetoencephalography data, Daume et al (2017) showed theta-alpha phase synchronization between inferior temporal cortex and frontopolar cortex during a visual WM task (71). Theta-theta phase synchronization between medial temporal lobe and PFC was observed in the context of visuospatial WM (72) and theta-related information flow between the hippocampus and the auditory cortex was reported for a verbal WM task (73). Most recently, Zhang et al. (2025) recorded human intracranial data during a visual search task that showed increased top-down medial PFC to mesial temporal lobe beta-gamma coherence, phase locking, and phase-amplitude coupling for fixation on a memory target versus fixation on a distractor (74). Note that visual stimuli were on-screen during that task period. They also found opposite changes in beta-gamma cross-frequency coupling metrics from mesial temporal lobe to medial PFC bottom-up. This shares themes with our LMFG-RMTG PBC result. While the authors did not explicitly quantify bursts, the effect they show seems to follow the concept of phase-burst coupling. To our knowledge, characterization of macroscale phase-burst dynamics between frontal and temporal areas during WM has not been reported. The PBC concept and metric could be used to study coupling both within- and across-other frontal and sensory regions not described here and provide insights into the interactions among potential WM nodes.

Our study has several limitations. Electrode locations were clinically determined, and thus there is an inherent spatial sampling bias and practical sampling limitation. These human subjects have epilepsy; however, we removed data derived from the seizure onset zones and trials with significant interictal activity from the analyses rendering clean data in that regard. We rely on subjects’ satisfactory performance in the task as well as the subjects having a relatively heterogenous anatomical distribution of seizure onset zone locations (see **Table 1, Table S1**) as grounds to assume that these subjects’ WM mechanisms on average resemble those of healthy subjects. Subjects also met a minimum score criteria on cognitive testing.

## Conclusions

Our findings highlight that bursting occurs in a frontotemporal network in humans and that such bursting may support memory maintenance. Furthermore, that burst-mediated network coupling may be a potential mechanism by which information is maintained in WM. Future work may focus on whether the burst code is an item-specific code (13, 14). Also, such coupling may extend to other brain areas, such as the hippocampus which is known to be important for both short- and long-term memory function (75). Delving deeper into burst features such as burst duration and their role in supporting such linkages may be another promising direction to advance our understanding of the human WM network.

## Materials and Methods

### Participants

This protocol was approved by the University of Utah Institutional Review Board and the University of Missouri Institutional Review Board. Participants were recruited from epilepsy patients with implanted electroencephalography or depth electrodes for long-term seizure monitoring. Inclusion criteria are a full-scale IQ greater than 70 or its equivalent based on cognitive testing, English speaking, >18 years old, and deemed clinically able to consent for themselves. Written consent was obtained from all participants by the research team, separate from the clinical team, and the patient was informed that study participation did not affect their clinical care in any way.

### Task design

The object WM task (**Figure 1a**) consisted of a fixation period (1000 ms) followed by the presentation of three consecutive visual stimuli (stimulus 1, 2, and 3; 500 ms each; shown as gray-shaded areas in figures) from one of three categories (faces, places or scenes). This sequence of task periods is referred to as the ‘trial scheme’. Items were presented in either left (−5.22 degrees offset, subtended 4.45 visual degrees), center (0 degrees offset, subtended 4.47 visual degrees), or right (5.22 degrees offset, subtended 4.45 visual degrees) position. After a delay period (4000 ms), subjects were shown a grid of 9 items and instructed to choose the stimuli shown during encoding by clicking on them with the mouse cursor. Subjects were given immediate feedback regarding which items were chosen correctly. Note that we did not consider item location as a condition for these analyses. To achieve a sufficient degree of dissimilarity between performance results, we defined ‘correct’ trials as those having all 3 items chosen correctly, and ‘incorrect’ trials as those with <2 correct items. Object stimuli were taken from the BOSS (Bank of Standardized Stimuli) database (76). Faces were taken from the FaceScrub database (77). Scenes were taken from the GAPED stimulus set (Geneva affective picture database) (78). Subjects in the Utah cohort were asked to maintain fixation during the trial period, but subjects in the Missouri cohort were not required to do so.

### Eye tracker

For the Utah cohort, eye position was sampled at 200 Hz using a camera mounted on glasses (Pupil Labs Core, Berlin). The eye tracker was calibrated at the start of the session and checked periodically. Deviations of greater than 10 degrees from central fixation aborted the trial. For the Missouri cohort, eye position was sampled at 150 Hz using a screen mounted camera (Gazepoint, Vancouver). Deviations of greater than 10 visual degrees from central fixation aborted the trial.

### Data recording and preprocessing

Electrode placement was determined entirely by clinical considerations. Electrode arrays were 4-12-channel linear arrays (10 mm spacing, either depth or subdural electrodes), typically with 8-12 arrays per patient. We chose a reference electrode positioned in the white matter that showed minimal electrophysiological signal.

All data were acquired using a 256-channel amplifier system (Blackrock Microsystems) at a sampling rate of 30 kHz, and then downsampled to 1 kHz. Trials with interictal discharges and artifact voltage deviations >1 mV were excluded. Channels with greater than 40% trials with significant artifact were removed. Channels within the clinical seizure-onset zone were removed. Finally, all data were notch-filtered at 60, 120, and 240 Hz, and then bipolar re-referenced and epoched into trials. Trials with interictal discharges (voltage fluctuations >300 mV above or below baseline) or with amplitude mean or variance greater than 3 SD of the ROI averages per subject (75) were removed.

### Detection of broadband-modulated channels

We computed broadband power (30–140 Hz) for each trial’s fixation and stimulus 1 period (5^th^ order Butterworth filter applied in the forward and reverse directions followed by squaring the absolute value of the Hilbert transform). We identified broadband-modulated channels as those with power during stimulus 1 that was 2 SD above or below the mean power during fixation. These were the only channels used in subsequent analyses.

### Burst extraction

We based our extraction algorithm on the procedure described in prior studies (9, 10). Oscillatory bursts in each frequency band (12-30 Hz for beta and 70-140 Hz for high gamma) were defined as time periods when the trial’s power exceeded 2 SD over the baseline power for at least 3 cycles (based on the band’s mean frequency). All power estimates were obtained with the procedure outlined above, and the baseline was computed by averaging power during the fixation period in a 10-trial window (current and nine previous trials). Each burst period was then refined by analyzing a 20-ms window around the burst’s bounds and extending the burst span for the duration that the power continuously exceeded 50% of the burst’s peak power.

### Burst rate computation and analysis

We represented each trial as a 0/1 vector with 1’s at indices corresponding to the time periods of detected bursts in 1 ms bins, over 0 to 6500 ms (separately for beta and high gamma bands). Two types of burst rates were calculated, (1) the trial-by-trial burst rate and (2) the 1/trial burst rate. The trial-by-trial burst rate is determined by calculating the number of bursts per second over a single trial. The 1/trial burst rate was calculated following (9) by averaging these 0/1 vectors over all trials, for each channel. Burst rate was baselined by subtracting the average rate during fixation. All statistical analyses performed on both trial-by-trial burst rate and 1/trial burst rates treated channel as the unit of observation.

### Phase-burst coupling

To assess the degree of high gamma burst coupling to beta phase, we adapted measures used in the theta-gamma single cell coupling analyses (41, 79). For each trial and channel pair, we computed the instantaneous beta phase trace for one channel and extracted high gamma bursts in the other channel. Phase was divided into 18 bins (−180 to 180 degrees in steps of 20 degrees). If the second channel had an ongoing burst at that time point, the corresponding phase bin was marked. The resulting distributions were averaged across trials for each channel pair. Delay-period distribution histograms reported were normalized to densities and were aggregated over the delay period from 3000 ms to 5500 ms.

The phase-burst coupling (PBC) measure is based on the modulation index from (41–43) computed in a sliding window over the trial (43). For each time point *t*, we constructed a bursts per phase distribution by aggregating bursts per phase counts over a 150 ms time window (from *t to t+150 samples)* and over trials. This procedure was performed to ensure that at least 200 beta cycles were present per window position (42) and that time windows were represented by the same number of beta cycles. We then calculated the relative difference between the entropy of the uniform distribution and the entropy of the instantaneous burst per phase distribution (ent_t_) using the following formula,

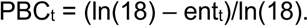

Note that in the case of within-ROI analyses, the phase and bursts were derived from a single channel to calculate a PBC for that channel. The PBC was then averaged over channels within an ROI.

### Statistical analyses

All metrics over time were tested using a cluster-based permutation test using 1000 permutations. We applied the cluster-based correction from (80). Permutation testing and cluster detection were performed using scipy (81), and clusters were identified only in the time dimension. This procedure was applied for testing significance of 1/trial burst rate, mutual information, PPL, PLV, and PBC. For dPLI, clusters were identified across both time and frequency domains.

Generalized linear models were estimated using MATLAB’s *fit_glme()* function with subject as a repeated measure. Pairwise comparisons and Tukey corrections were performed by using MATLAB’s *anovan()* and *multcompare()* functions.

Rayleigh tests were performed using the astropy package (82). All correlation estimates refer to Spearman’s rank correlation coefficient.

### Additional analyses

Additional analyses are available in *Supplementary Materials*.

## Data and code availability

Data (doi:10.18112/openneuro.ds006136.v1.0.0) and code (Python, MATLAB in a GitHub repository) used for analyses are available.

## Acknowledgments

This work was supported by a K12 grant from the Neurosurgery Career Development Award, parent award from the NINDS as well as a Neurosurgery Research Education Fund award, NIH/NINDS career development award to JDR (K23 NS114178), and NIMH grant MH122023 to SSN.

We would like to thank Dr. Nelson Cowan and Dr. Drew B. Headley for their helpful comments.

## References

1. C. Constantinidis, et al., Persistent spiking activity underlies working memory. J Neurosci 38, 7020–7028 (2018).

2. M. Lundqvist, P. Herman, E. K. Miller, Working memory: delay activity, yes! Persistent activity? Maybe not. J Neurosci 38, 7013–7019 (2018).

3. C. E. Curtis, T. C. Sprague, Persistent activity during working memory from front to back. Front Neural Circuits 15, 696060 (2021).

4. J. M. Fuster, G. E. Alexander, Neuron activity related to short-term memory. Science 173, 652–654 (1971).

5. S. Funahashi, C. J. Bruce, P. S. Goldman-Rakic, Mnemonic coding of visual space in the monkey’s dorsolateral prefrontal cortex. J Neurophysiol 61, 331–349 (1989).

6. S. Kojima, P. S. Goldman-Rakic, Delay-related activity of prefrontal neurons in rhesus monkeys performing delayed response. Brain Res 248, 43–50 (1982).

7. C. Constantinidis, M. N. Franowicz, P. S. Goldman-Rakic, The sensory nature of mnemonic representation in the primate prefrontal cortex. Nat Neurosci 4, 311–316 (2001).

8. J. Kamiński, et al., Persistently active neurons in human medial frontal and medial temporal lobe support working memory. Nat Neurosci 20, 590–601 (2017).

9. M. Lundqvist, et al., Gamma and beta bursts underlie working memory. Neuron 90, 152– 164 (2016).

10. M. Lundqvist, P. Herman, M. R. Warden, S. L. Brincat, E. K. Miller, Gamma and beta bursts during working memory readout suggest roles in its volitional control. Nat Commun 9, 394 (2018).

11. E. K. Miller, M. Lundqvist, A. M. Bastos, Working memory 2.0. Neuron 100, 463–475 (2018).

12. M. Lundqvist, et al., Working memory control dynamics follow principles of spatial computing. Nat Commun 14, 1429 (2023).

13. W. Xie, et al., Neuronal sequences in population bursts encode information in human cortex. Nature 1–8 (2024).

14. M. F. Panichello, et al., Intermittent rate coding and cue-specific ensembles support working memory. Nature 1–8 (2024).

15. M. Balconi, Dorsolateral prefrontal cortex, working memory and episodic memory processes: insight through transcranial magnetic stimulation techniques. Neurosci Bull 29, 381–389 (2013).

16. R. D. Webler, et al., DLPFC stimulation alters working memory related activations and performance: An interleaved TMS-fMRI study. Brain Stimul 15, 823–832 (2022).

17. Y. Miyashita, H. S. Chang, Neuronal correlate of pictorial short-term memory in the primate temporal cortex. Nature 331, 68–70 (1988).

18. L. Chelazzi, J. Duncan, E. K. Miller, R. Desimone, Responses of neurons in inferior temporal cortex during memory-guided visual search. J Neurophysiol 80, 2918–2940 (1998).

19. D. Zaksas, J. W. Bisley, T. Pasternak, Motion information is spatially localized in a visual working-memory task. J Neurophysiol 86, 912–921 (2001).

20. T. Pasternak, M. W. Greenlee, Working memory in primate sensory systems. Nat Rev Neurosci 6, 97–107 (2005).

21. B. R. Postle, Working memory as an emergent property of the mind and brain. Neuroscience 139, 23–38 (2006).

22. P. Jolicœur, C. Lefebvre, J. Martinez-Trujillo, Mechanisms of Sensory Working Memory: Attention and Performance XXV (Academic Press, 2015).

23. N. S. Rose, et al., Reactivation of latent working memories with transcranial magnetic stimulation. Science 354, 1136–1139 (2016).

24. K. K. Sreenivasan, M. D’Esposito, The what, where and how of delay activity. Nat Rev Neurosci 20, 466–481 (2019).

25. B. Pesaran, J. S. Pezaris, M. Sahani, P. P. Mitra, R. A. Andersen, Temporal structure in neuronal activity during working memory in macaque parietal cortex. Nat Neurosci 5, 805– 811 (2002).

26. E. K. Miller, L. Li, R. Desimone, A neural mechanism for working and recognition memory in inferior temporal cortex. Science 254, 1377–1379 (1991).

27. E. K. Miller, L. Li, R. Desimone, Activity of neurons in anterior inferior temporal cortex during a short-term memory task. J Neurosci 13, 1460–1478 (1993).

28. M. Dake, C. E. Curtis, Perturbing human V1 degrades the fidelity of visual working memory. Nat Commun 16, 2675 (2025).

29. E. Rezayat, K. Clark, M.-R. A. Dehaqani, B. Noudoost, Dependence of working memory on coordinated activity across brain areas. Front Syst Neurosci 15, 787316 (2022).

30. P. Comeaux, K. Clark, B. Noudoost, A recruitment through coherence theory of working memory. Prog Neurobiol 228, 102491 (2023).

31. E. Rezayat, et al., Frontotemporal coordination predicts working memory performance and its local neural signatures. Nat Commun 12, 1103 (2021).

32. R. G. Briggs, et al., The unique fiber anatomy of middle temporal gyrus default mode connectivity. Oper Neurosurg 21, E8–E14 (2021).

33. M. Petrides, On the evolution of polysensory superior temporal sulcus and middle temporal gyrus: A key component of the semantic system in the human brain. J Comp Neurol 531, 1987–1995 (2023).

34. Y. Norman, et al., Hippocampal sharp-wave ripples linked to visual episodic recollection in humans. Science 365, eaax1030 (2019).

35. T. P. Zanto, M. T. Rubens, A. Thangavel, A. Gazzaley, Causal role of the prefrontal cortex in top-down modulation of visual processing and working memory. Nat Neurosci 14, 656– 661 (2011).

36. B. Noudoost, T. Moore, Control of visual cortical signals by prefrontal dopamine. Nature 474, 372–375 (2011).

37. R. F. Helfrich, R. T. Knight, Oscillatory dynamics of prefrontal cognitive control. Trends Cogn Sci 20, 916–930 (2016).

38. A. Dubey, D. A. Markowitz, B. Pesaran, Top-down control of exogenous attentional selection is mediated by beta coherence in prefrontal cortex. Neuron 111, 3321–3334 (2023).

39. L. G. Ungerleider, S. M. Courtney, J. V Haxby, A neural system for human visual working memory. Proc Natl Acad Sci USA 95, 883–890 (1998).

40. C. J. Stam, E. C. W. van Straaten, Go with the flow: use of a directed phase lag index (dPLI) to characterize patterns of phase relations in a large-scale model of brain dynamics. Neuroimage 62, 1415–1428 (2012).

41. A. B. L. Tort, R. W. Komorowski, J. R. Manns, N. J. Kopell, H. Eichenbaum, Theta–gamma coupling increases during the learning of item–context associations. Proc Natl Acad Sci USA 106, 20942–20947 (2009).

42. A. B. L. Tort, R. Komorowski, H. Eichenbaum, N. Kopell, Measuring phase-amplitude coupling between neuronal oscillations of different frequencies. J Neurophysiol 104, 1195– 1210 (2010).

43. S. Samiee, S. Baillet, Time-resolved phase-amplitude coupling in neural oscillations. Neuroimage 159, 270–279 (2017).

44. E. D. Gribkova, B. A. Ibrahim, D. A. Llano, A novel mutual information estimator to measure spike train correlations in a model thalamocortical network. J Neurophysiol 120, 2730–2744 (2018).

45. C. Tallon-Baudry, O. Bertrand, C. Fischer, Oscillatory synchrony between human extrastriate areas during visual short-term memory maintenance. J Neurosci 21, RC177 (2001).

46. J. M. Palva, S. Monto, S. Kulashekhar, S. Palva, Neuronal synchrony reveals working memory networks and predicts individual memory capacity. Proc Natl Acad Sci USA 107, 7580–7585 (2010).

47. R. Desimone, J. Duncan, Neural mechanisms of selective visual attention. Annu Rev Neurosci 18, 193–222 (1995).

48. A. Mikami, S. Ito, K. Kubota, Visual response properties of dorsolateral prefrontal neurons during visual fixation task. J Neurophysiol 47, 593–605 (1982).

49. E. A. Buffalo, P. Fries, R. Landman, T. J. Buschman, R. Desimone, Laminar differences in gamma and alpha coherence in the ventral stream. Proc Natl Acad Sci USA 108, 11262– 11267 (2011).

50. A. M. Bastos, R. Loonis, S. Kornblith, M. Lundqvist, E. K. Miller, Laminar recordings in frontal cortex suggest distinct layers for maintenance and control of working memory. Proc Natl Acad Sci USA 115, 1117–1122 (2018).

51. M. W. Howard, et al., Gamma oscillations correlate with working memory load in humans. Cereb Cortex 13, 1369–1374 (2003).

52. M. Lundqvist, E. K. Miller, J. Nordmark, J. Liljefors, P. Herman, Beta: bursts of cognition. Trends Cogn Sci (2024).

53. C. de Hemptinne, P. Lefevre, M. Missal, Neuronal bases of directional expectation and anticipatory pursuit. J Neurosci 28, 4298–4310 (2008).

54. S. Picazio, et al., Prefrontal control over motor cortex cycles at beta frequency during movement inhibition. Curr Biol 24, 2940–2945 (2014).

55. S. Little, J. Bonaiuto, G. Barnes, S. Bestmann, Human motor cortical beta bursts relate to movement planning and response errors. PLoS Biol 17, e3000479 (2019).

56. R. Lofredi, et al., Beta bursts during continuous movements accompany the velocity decrement in Parkinson’s disease patients. Neurobiol Dis 127, 462–471 (2019).

57. D. L. Paulo, et al., Corticostriatal beta oscillation changes associated with cognitive function in Parkinson’s disease. Brain 146, 3662–3675 (2023).

58. A. K. Barbey, M. Koenigs, J. Grafman, Dorsolateral prefrontal contributions to human working memory. Cortex 49, 1195–1205 (2013).

59. D. A. Cohen, E. M. Robertson, Preventing interference between different memory tasks. Nat Neurosci 14, 953–955 (2011).

60. E. M. Robertson, New insights in human memory interference and consolidation. Curr Biol 22, R66–R71 (2012).

61. J. Liljefors, et al., Distinct functions for beta and alpha bursts in gating of human working memory. Nat Commun 15, 8950 (2024).

62. M. T. Kucewicz, et al., Dissecting gamma frequency activity during human memory processing. Brain 140, 1337–1350 (2017).

63. M. Silva, et al., Movie-watching evokes ripple-like activity within events and at event boundaries. Nat Commun 16, 5647 (2025).

64. R. F. Salazar, N. M. Dotson, S. L. Bressler, C. M. Gray, Content-specific fronto-parietal synchronization during visual working memory. Science 338, 1097–1100 (2012).

65. A. B. L. Tort, et al., Dynamic cross-frequency couplings of local field potential oscillations in rat striatum and hippocampus during performance of a T-maze task. Proc Natl Acad Sci USA 105, 20517–20522 (2008).

66. N. Suthana, et al., Memory enhancement and deep-brain stimulation of the entorhinal area. N Engl J Med 366, 502–510 (2012).

67. I. Alekseichuk, Z. Turi, G. A. De Lara, A. Antal, W. Paulus, Spatial working memory in humans depends on theta and high gamma synchronization in the prefrontal cortex. Curr Biol 26, 1513–1521 (2016).

68. C. Inman, et al., Direct Electrical Stimulation of the Amygdala Enhances Event-Specific Declarative Memory in Humans. Proc Natl Acad Sci USA 95 (2017).

69. Y. Ezzyat, et al., Closed-loop stimulation of temporal cortex rescues functional networks and improves memory. Nat Commun 9, 365 (2018).

70. J. Daume, et al., Control of working memory by phase–amplitude coupling of human hippocampal neurons. Nature 629, 393–401 (2024).

71. J. Daume, T. Gruber, A. K. Engel, U. Friese, Phase-amplitude coupling and long-range phase synchronization reveal frontotemporal interactions during visual working memory. J Neurosci 37, 313–322 (2017).

72. E. L. Johnson, et al., Dynamic frontotemporal systems process space and time in working memory. PLoS Biol 16, e2004274 (2018).

73. V. Dimakopoulos, P. Mégevand, L. H. Stieglitz, L. Imbach, J. Sarnthein, Information flows from hippocampus to auditory cortex during replay of verbal working memory items. Elife 11, e78677 (2022).

74. J. Zhang, J. Xia, H. Zhou, S. Wang, Gamma synchronization between the medial temporal lobe and medial frontal cortex for goal-directed visual attention in humans. Cell Rep 44 (2025).

75. W. Xie, et al., The medial temporal lobe supports the quality of visual short-term memory representation. Nat Hum Behav 7, 627–641 (2023).

76. M. B. Brodeur, K. Guérard, M. Bouras, Bank of Standardized Stimuli (BOSS) phase II: 930 new normative photos. PLoS One 9, e106953 (2014).

77. H.-W. Ng, S. Winkler, A data-driven approach to cleaning large face datasets in 2014 IEEE International Conference on Image Processing (ICIP), (IEEE, 2014), pp. 343–347.

78. E. S. Dan-Glauser, K. R. Scherer, The Geneva affective picture database (GAPED): a new 730-picture database focusing on valence and normative significance. Behav Res Methods 43, 468–477 (2011).

79. A. Sirota, et al., Entrainment of neocortical neurons and gamma oscillations by the hippocampal theta rhythm. Neuron 60, 683–697 (2008).

80. E. Maris, R. Oostenveld, Nonparametric statistical testing of EEG-and MEG-data. J Neurosci Methods 164, 177–190 (2007).

81. P. Virtanen, et al., SciPy 1.0: fundamental algorithms for scientific computing in Python. Nat Methods 17, 261–272 (2020).

82. A. M. Price-Whelan, et al., The Astropy Project: sustaining and growing a community-oriented open-source project and the latest major release (v5. 0) of the core package. Astrophys J 935, 167 (2022).

