## Supplementary Information for "Frontotemporal bursting supports human working memory"

### Supporting Information for
Frontotemporal bursting supports human working memory

Vladimir Omelyusik[1], Tyler S. Davis[2], Satish S. Nair*[1], Behrad Noudoost[3], Patrick D. Hackett[4], Elliot H. Smith[2], Shervin Rahimpour[2], John D. Rolston[2,5], Bornali Kundu*[1,2,4]

1. Department of Electrical Engineering and Computer Science, University of Missouri, Columbia, MO, 65211. 2. Department of Neurosurgery, University of Utah, Salt Lake City, UT 84132. 3. Department of Ophthalmology, University of Utah, Salt Lake City, UT 84132. 4. Department of Neurosurgery, University of Missouri, Columbia, MO, 65211. 5. Department of Neurosurgery, Mass General Brigham, Harvard Medical School, Boston, MA 02115.

* Bornali Kundu
One Hospital Drive
MC 314
Columbia, MO 65213

* Satish S. Nair (co-corresponding author)
S 6th Street
229 Naka Hall
Columbia, MO 65213


**This PDF file includes:**

Supporting text
SI Methods and Results
Figures S1 to S16
Tables S1 to S2
SI References

#### SI Methods and Results

##### Burst rate versus number correct

In the main text we define 'incorrect' trials as those having 0 or 1 item chosen correctly during the search period. To assess whether burst rate scales with the number of items correct, we repeated the performance analysis with 'incorrect' trials being defined as those having 0, 1, or 2 (i.e., <3) items chosen correctly (**Figure S16**). We found evidence that burst rates scale with number correct.

##### PBC versus WM performance

The PBC measure is sensitive to the number of cycles over which it is computed. Since the number of incorrect trials (and hence the number of cycles per window position) was lower compared to the number of correct trials (the mean number of correct trials was 80.35 and the mean number of incorrect trials was 29.3 per channel pair), we constructed an approximate trial-balanced estimate of PBC for incorrect trials. For each channel pair, we shuffled trial order and then partitioned correct trials into *n* partitions containing the same number of trials as there were incorrect trials. We then computed PBC for each Correct partition (~5 partitions). We then computed a single PBC value over all incorrect trials and upsampled *n* times to match the number of Correct partitions. These PBC values for correct and incorrect trials were aggregated across channel pairs within a region pair (e.g. RMTG and LMFG) and tested for significance using a 2-way ANOVA (**Figure S9a**) and a cluster-based permutation test (**Figure S9b**). We found elevated delay period beta – high gamma PBC for correct vs incorrect trials within the LMFG-RMTG pair. For the RMTG-LMFG pair, PBC during incorrect trials was significantly elevated during the stimulus 1 period of encoding.

##### Local and long-range PBC

To explore the PBC measure, we computed the beta-high gamma PBC between LMFG and RMTG (**Figure 5c**), LMFG and LMTG (**Figure S10a**), and at individual channels within these regions (**Figure S10b**) for correct trials. We found sustained elevated PBC only in the first case ($p < 0.05$, cluster test). Since PBC may be function specific, and thus not elevated uniformly across all available pairs, we then selected the top 20% of channel pairs (between regions) or channels (within regions) with the highest delay period PBC and compared PBC between correct and incorrect trials in a trial-balanced way described above. Notably, there was a significant difference between correct and incorrect trials ($p < 0.05$, cluster test) in the beta – high gamma PBC between RMTG and LMFG pairs (**Figure S11a**), as opposed to PBC computed over all pairs (**Figure S9b**). Periods of significant differences were found within each region as well (**Figure S11b**). Future research should explore appropriate selection of channel pairs for PBC analysis.

##### Mutual information

We identified all possible pairs of LMFG and RMTG channels for each patient and computed mutual information between high gamma bursts during 100-900 ms of fixation and 3 sub-periods of the delay period (2600-3400 ms, 4000-4800 ms, 5200-6000 ms) of each trial. We used a mutual information estimate for spike trains and adapted it to bursts (1). For every window position, we had two "burst trains" represented by 0/1 vectors, one from each channel in the LMFG-RMTG pair. We then determined the number of bursts that co-occurred per channel pair. Mutual information was computed as the sum of entropies of marginal distributions minus the entropy of the joint distribution. The estimates were then averaged across all trials per channel pair. All statistical analyses treated channel pairs as a unit of observation.

##### Phase locking

Beta-beta phase locking was estimated via PPL (2). Similar to mutual information, PPL and PLV measures were estimated per trial in a 150-ms running window. Per-trial measures were then averaged across trials and region pairs.

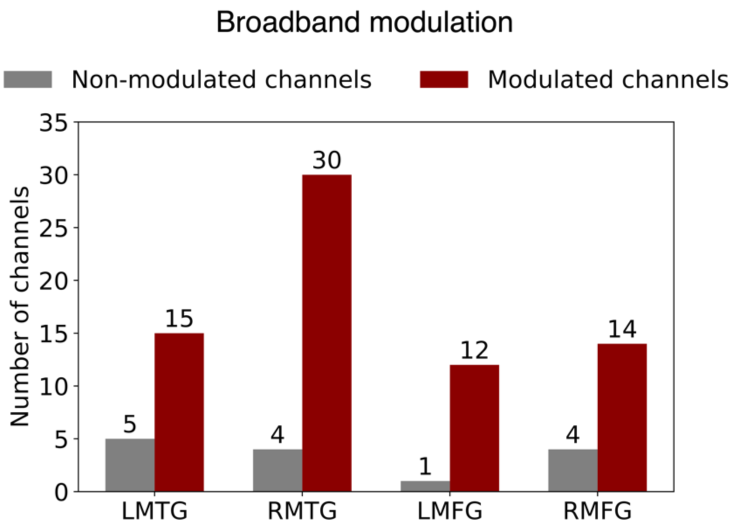


**Fig. S1. Number of broadband-modulated channels across ROIs |** These channels showed significant broadband (30-140 Hz) activity during the encoding period of stimulus 1.

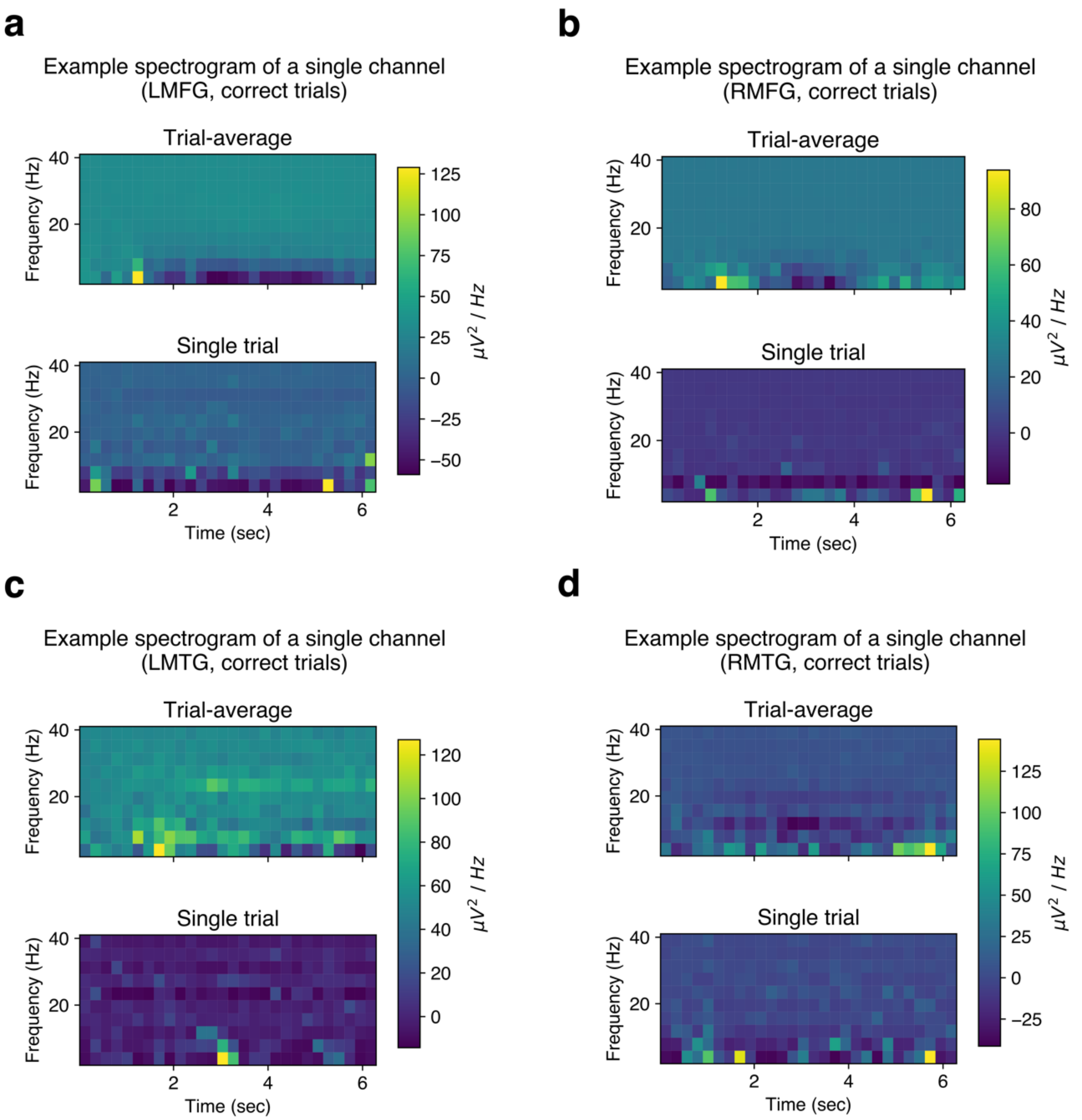


**Fig. S2. Examples of trial-averaged and single-trial activity in the 1-40 Hz frequency band across ROIs | a-d.** Similar to the high gamma frequency band, power in the beta (12-30 Hz) band was elevated over shorter time periods in single trials compared to trial-averages.

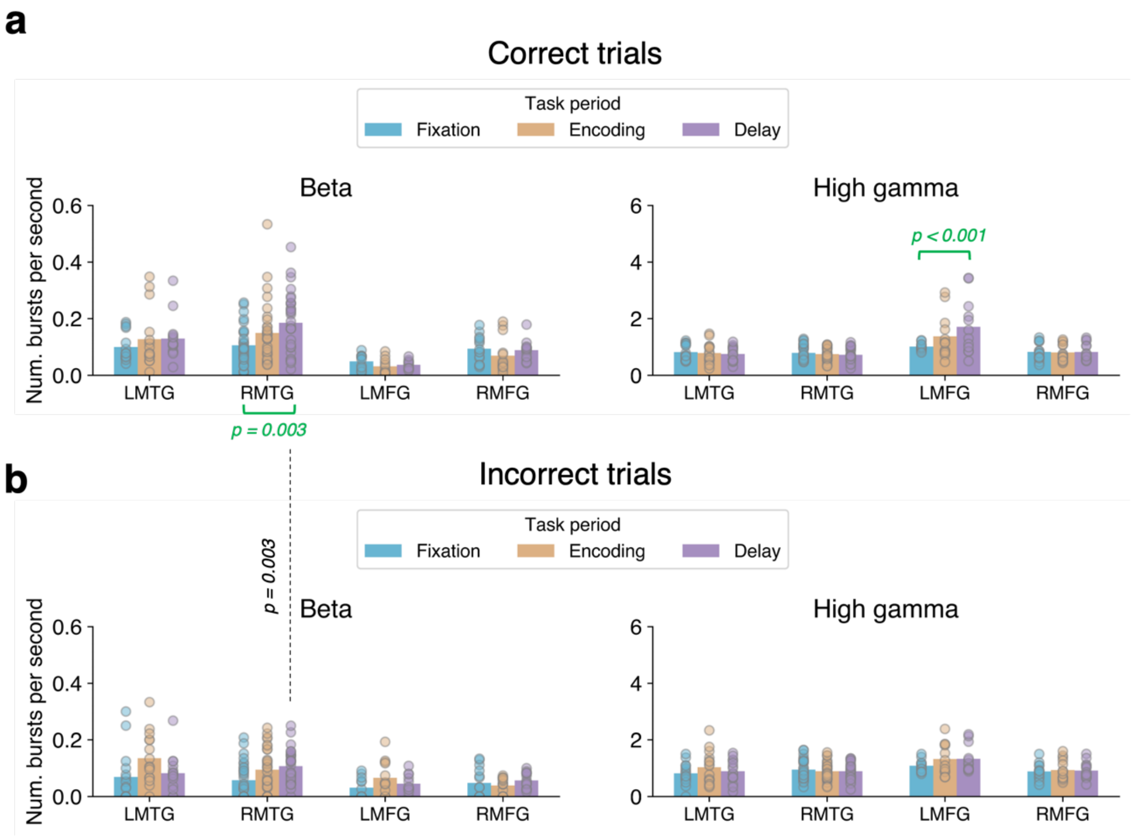


**Fig. S3. Trial-by-trial beta and high gamma burst rate (in bursts per second) for each task period and ROI |** Note this measure is different from the trial-average (i.e. 1/trial) burst rate referred to in other figures. The per second rate is calculated over the time of the trial (i.e. trial-by-trial). Beta and high gamma bursts were detected in all regions and task periods, for both correct (**a**) and incorrect (**b**) trials. Bar plots show post hoc pairwise tests with Tukey correction. 2-way GLME for each ROI and band. **LMFG, beta**: significant main effect of period ($F$ = 3.15, $DF1$ = 2, $DF2$ = 66, $p$ = 0.0495), no significant main effect of performance, significant performance x period interaction ($F$ = 3.47, $DF1$ = 2, $DF2$ = 66, $p$ = 0.04). **LMFG, high gamma**: no significant effects of period, performance or interaction. **RMFG, beta**: significant main effect of performance ($F$ = 9.25, $DF1$ = 1, $DF2$ = 78, $p$ = 0.003), no other significant effects. **RMFG, high gamma**: no significant effects. **RMTG, beta**: significant main effect of period ($F$ = 3.72, $DF1$ = 2, $DF2$ = 174, $p$ = 0.03) and performance ($F$ = 6.57, $DF1$ = 1, $DF2$ = 174, $p$ = 0.01), no significant interaction. **RMTG, high gamma**: significant main effect of performance ($F$ = 6.14, $DF1$ = 1, $DF2$ = 174, $p$ = 0.01), no significant effects of period or interaction. **LMTG, beta**: significant main effect of period ($F$ = 3.25, $DF1$ = 2, $DF2$ = 84, $p$ = 0.04), no other significant effects. **LMTG, high gamma**: no significant effects found.

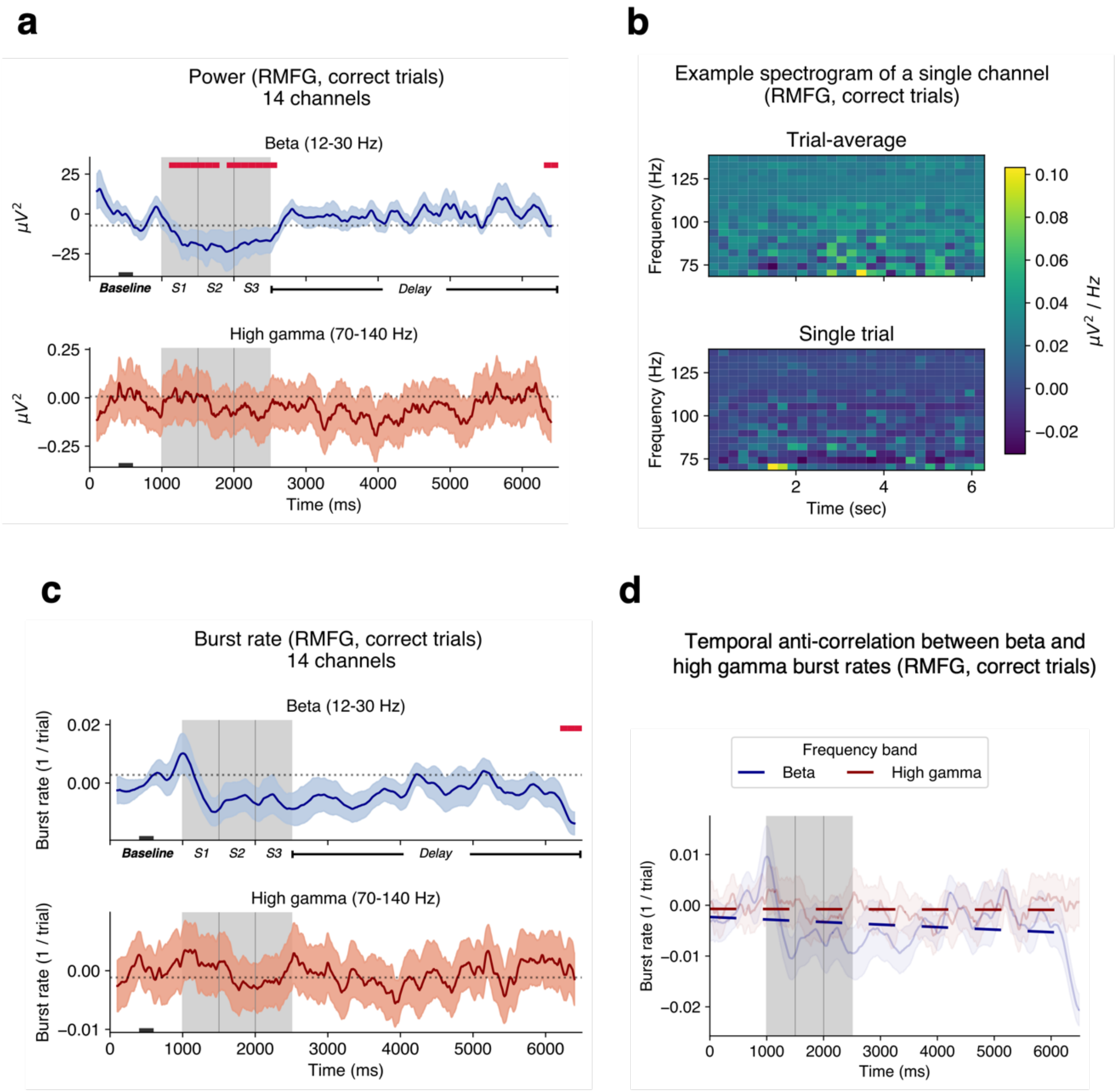


**Fig. S4. Activity in RMFG (correct trials, 14 channels, 4 subjects) | a.** Beta and high gamma power averaged over channels. Beta power decreased during encoding compared to the baseline. **b.** An example of trial-averaged and single-trial spectrograms in the high gamma frequency band (70-140 Hz). Periods of elevated power are longer in the trial-averaged spectrogram and shorter in the single-trial spectrogram. **c.** Beta and high gamma burst rates. No significant differences with fixation were found for the majority of the task duration. **d.** Anticorrelation between beta and high gamma burst rate over the trial (average $r = -0.03$ with $p < 0.05$ for 12 / 14 channels). In panels a and c, shaded regions correspond to ±1 SEM. Crimson lines indicate significant differences from the mean of 400-600 ms of fixation (black rectangle at the bottom; $p < 0.05$, cluster test; 200 ms running window, 50% overlap). A 200 ms smoothing window was used for visualization but not for testing.

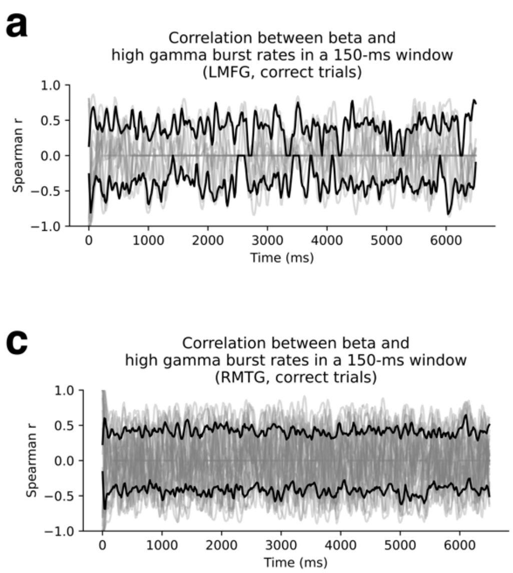


**Fig. S5. High gamma-beta burst rate anti-correlation is not observed in short time windows (LMFG: 12 channels, RMFG: 14 channels, RMTG: 30 channels, LMTG: 15 channels) | a-d.** Spearman rank correlations between high gamma and beta burst rates in 150-ms moving windows with 90% overlap. Gray lines show correlation traces in individual channels; black lines show averages across channels with positive and negative correlations.

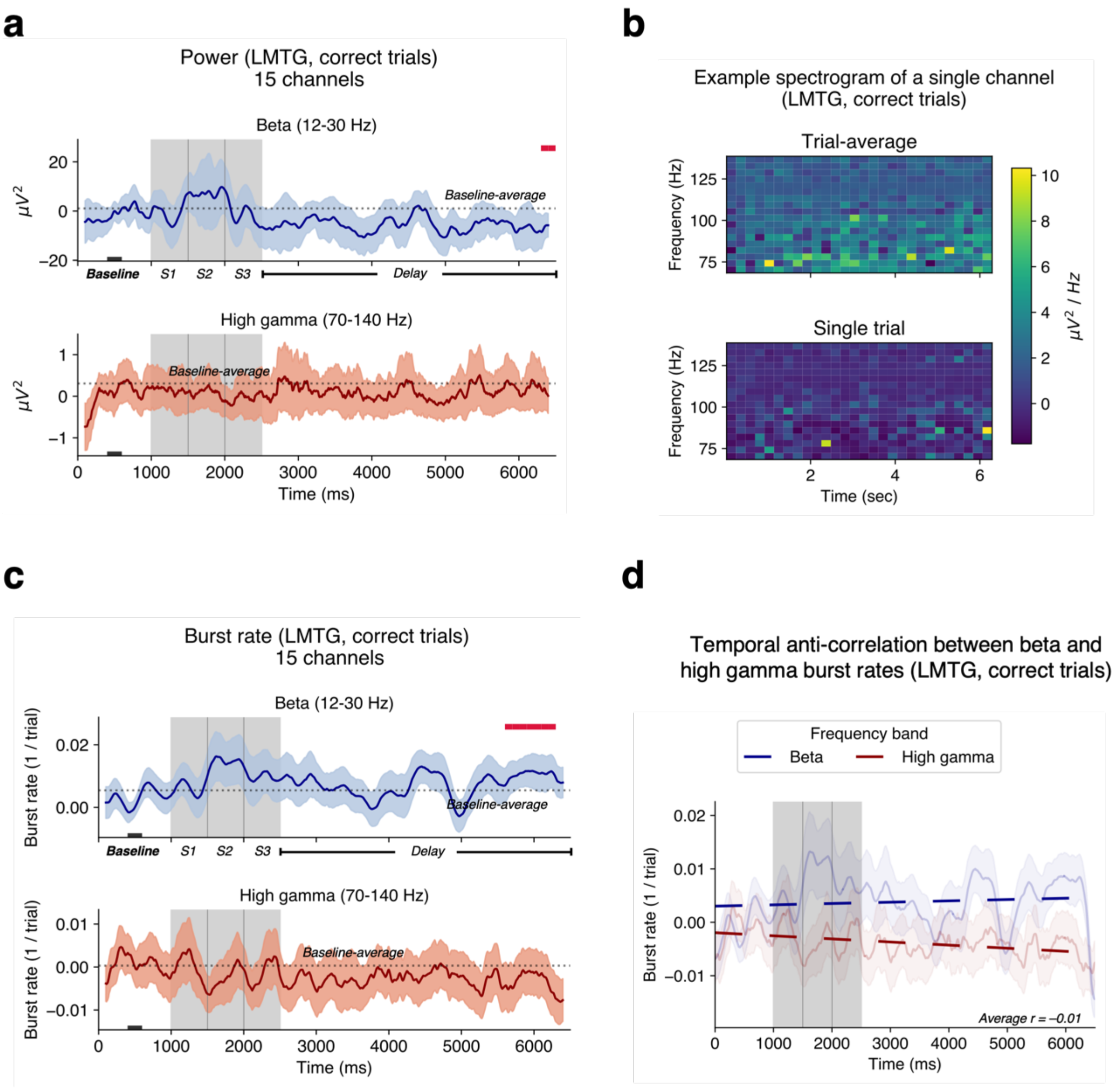


**Fig. S6. Activity in LMTG (correct trials, 15 channels, 5 subjects) | a.** Beta and high gamma power averaged over channels. Power in either band was not significantly different from baseline. **b.** An example of trial-averaged and single-trial spectrograms in the high gamma frequency band (70-140 Hz). Single trials show bursts of power. **c.** 1/trial beta and high gamma burst rates. Beta burst rate was elevated compared to the baseline at the end of the delay period. **d.** Anticorrelation of high gamma and beta burst rates over the trial (average $r$ = -0.01 with $ps$ < 0.05 for 12 / 15 channels). In panels **a** and **c**, shaded regions correspond to ±1 SEM. Crimson lines indicate significant differences from the mean of 400-600 ms of fixation (black rectangle at the bottom; $p$ < 0.05, cluster test; 200 ms running window, 50% overlap). A 200 ms smoothing window was used for visualization but not for testing.

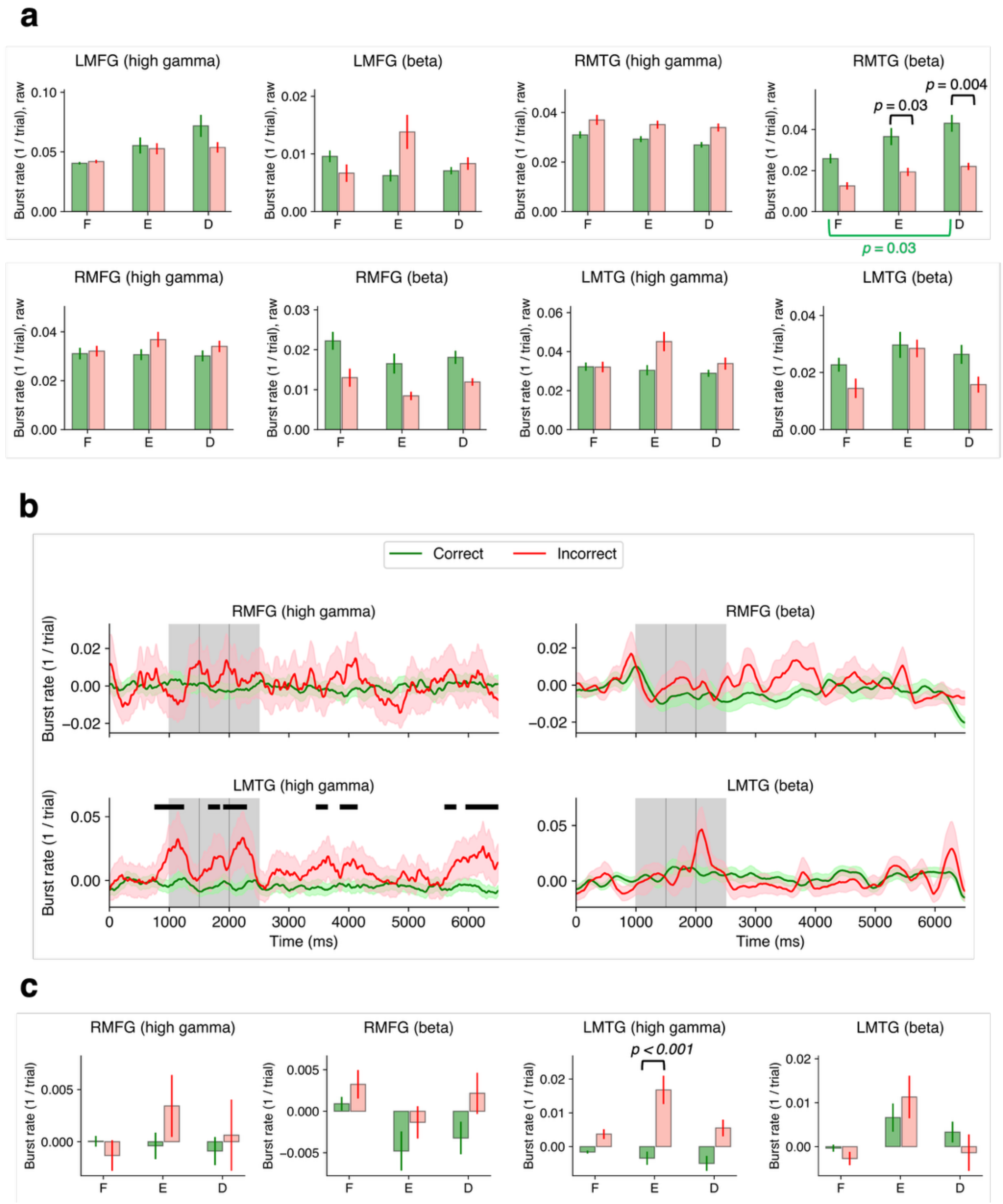


**Fig. S7. Burst rates in RMFG and LMTG for correct and incorrect trials | a.** Bar plots of raw 1/trial burst rates for correct (green) and incorrect (red) trials. For each region, 2-way GLME and post hoc pairwise tests with Tukey correction were done. **LMFG**: no significant interaction effect or main effect of period or main effect of performance on either raw beta or raw high gamma burst rates. **RMTG**: significant main effect of performance on raw high gamma ($F$ = 5.01, $DF1$ = 1, $DF2$ = 174, $p$ = 0.03) and beta ($F$ = 7.02, $DF1$ = 1, $DF2$ = 174, $p$ = 0.009) burst rates; no other significant effects. **RMFG**: significant main effect of performance on raw beta burst rate ($F$ = 7.53, $DF1$ = 1, $DF2$ = 78, $p$ = 0.008); no other significant effects. **LMTG**: significant main effect of period on raw high gamma ($F$ = 5.36, $DF1$ = 2, $DF2$ = 84, $p$ = 0.006) and beta ($F$ = 3.35, $DF1$ = 2, $DF2$ = 84, $p$ = 0.04) burst rates; no other significant effects. **b.** Temporal dynamics of baseline-subtracted 1/trial burst rates for correct and incorrect trials. Shaded regions represent ±1 SEM. Black lines correspond to periods of significant difference between correct and incorrect averages over channels ($p$ < 0.05, cluster test; 200 ms running window, 75% overlap). The baseline period was

200-800 ms from the start of the trial. A 200 ms smoothing window was used for visualization but not for testing. **c.** Bar plots of baseline-subtracted burst rates during 3 periods and 2 performance conditions. For each region, 2-way GLME and post hoc pairwise tests with Tukey correction were done. **RMFG**: no significant interaction effect or main effect of period or main effect of performance on either raw beta or raw high gamma burst rates. **LMTG**: main effect of period ($F$ = 6.05, $DF1$ = 2, $DF2$ = 84, $p$ = 0.003) and significant period x performance interaction ($F$ = 3.76, $DF1$ = 2, $DF2$ = 84, $p$ = 0.03) on high gamma burst rate and main effect of period ($F$ = 3.4, $DF1$ = 2, $DF2$ = 84, $p$ = 0.04) on beta burst rate; no other significant effects. Error bars represent ±1 SEM.

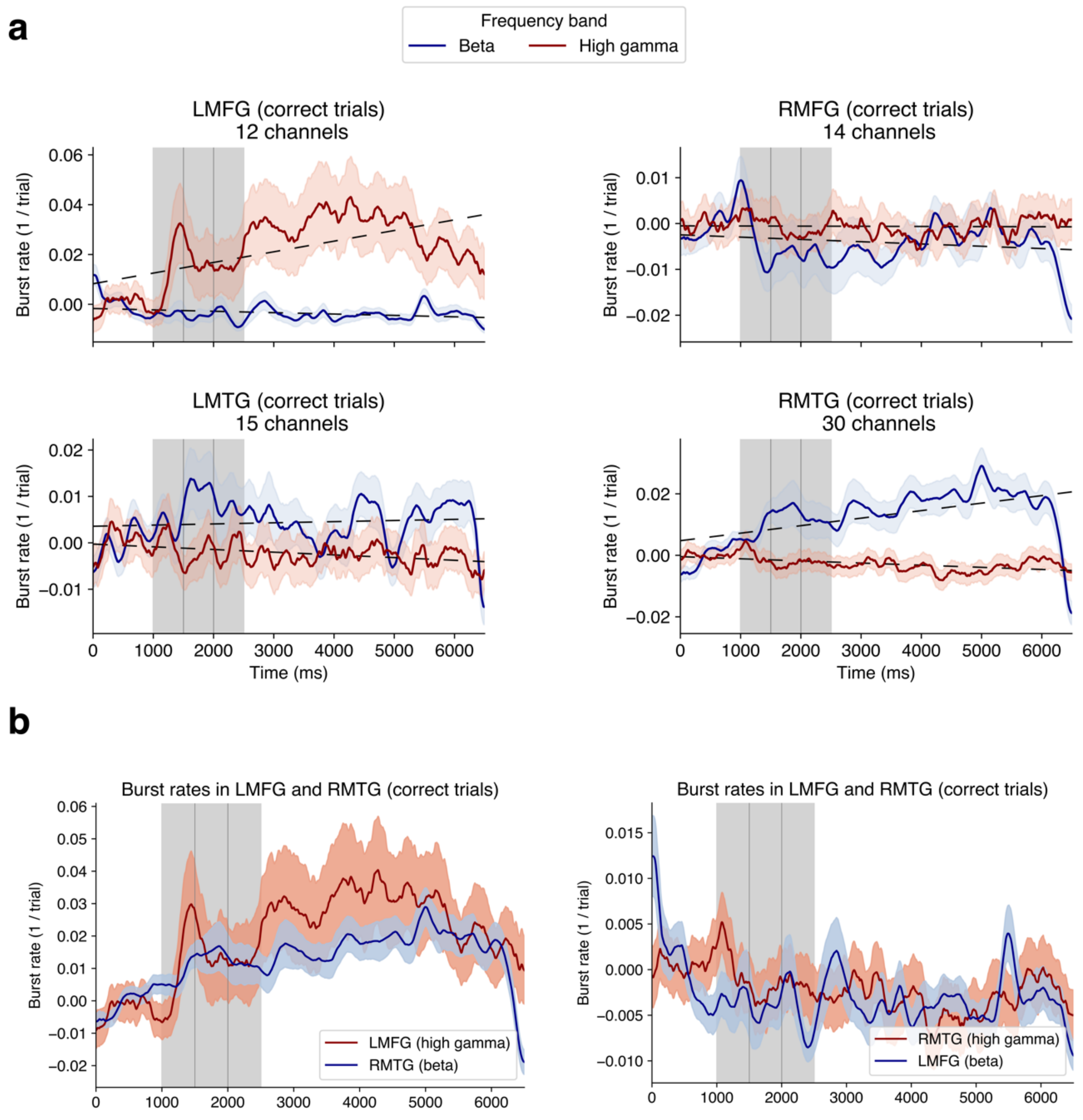


**Fig. S8. High gamma and beta burst rates are anticorrelated (correct trials) | a.** High gamma and beta burst rate dynamics in all ROIs. There was an increase in high gamma bursting in frontal areas and an increase in beta bursting in temporal areas. Shaded areas correspond to ±1 SEM, a 200 ms smoothing window was used for visualization. **b.** High gamma – beta burst rates in LMFG and RMTG follow similar temporal profiles.

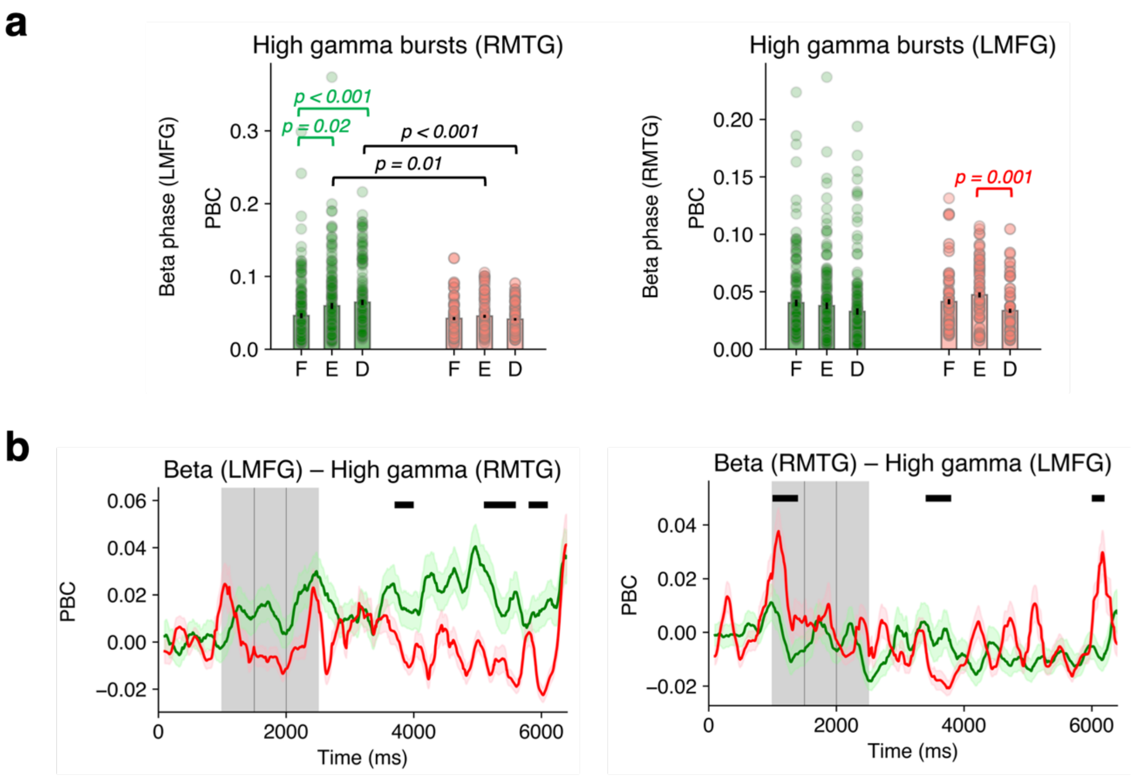


**Fig. S9. Phase-burst coupling (PBC) between LMFG and RMTG (42 channel pairs within LMFG, data from 3 subjects; 66 pairs between LMFG and RMTG, data from 2 subjects; 166 pairs within RMTG, data from 5 subjects). |** Temporal dynamics of PBC with matched numbers of correct and incorrect trials (see **SI Methods and Results**). **a.** Bar plots of PBC values without baseline subtraction. PBC was significantly elevated during the delay period compared to fixation in the LMFG-RMTG pair for correct trials compared to incorrect trials (post hoc pairwise tests, Tukey correction). Error bars correspond to ±1 SEM. **b.** Black lines indicate significant differences between correct and incorrect trials ($p < 0.05$, cluster test; 200 ms running window, 50% overlap). A 200 ms smoothing window was used for visualization but not for testing. Shaded regions represent ±1 SEM.

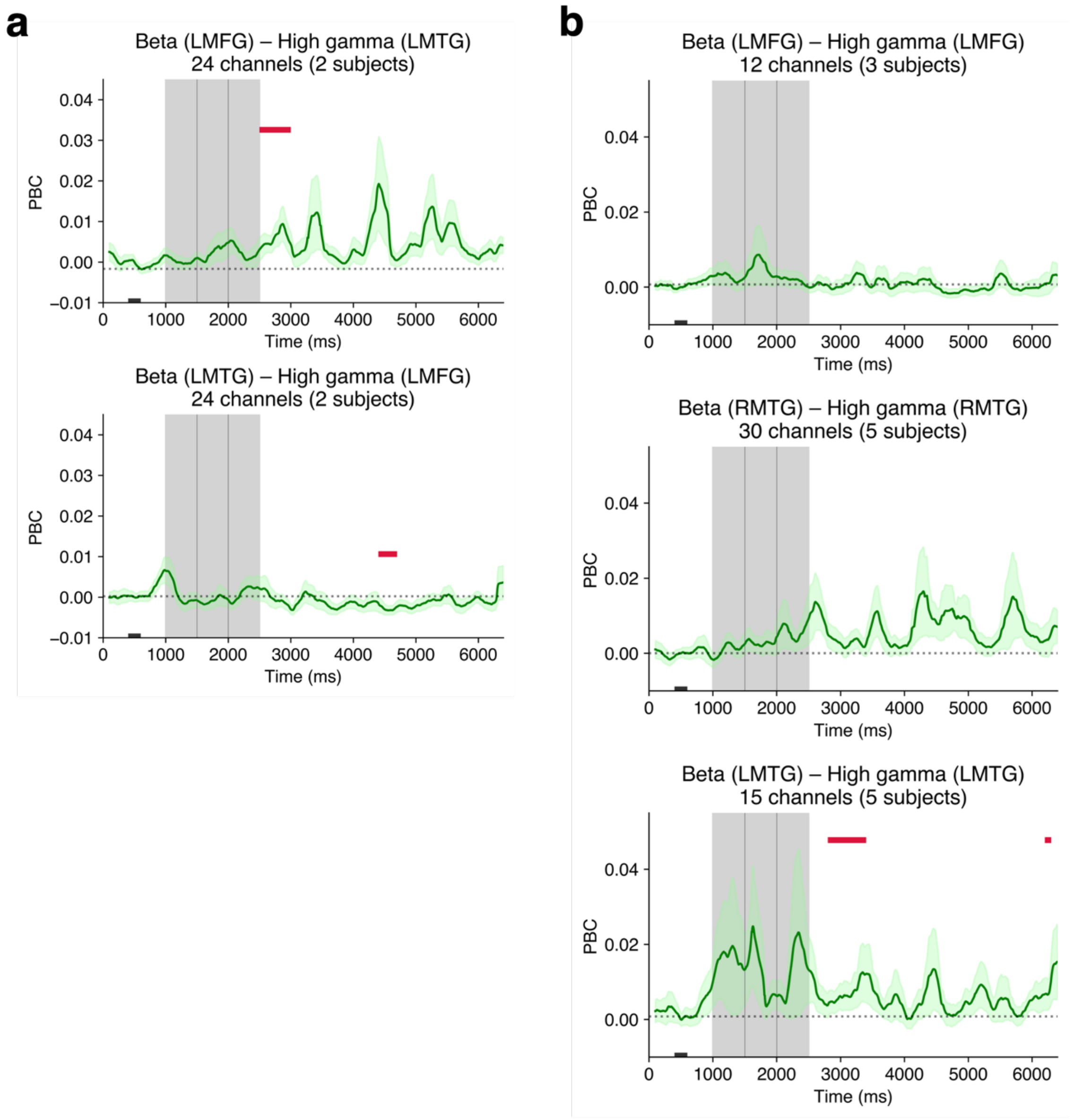


**Figure S10. PBC within and between frontal and temporal ROIs. | a.** Beta-high gamma PBC between LMFG and LMTG channels. **b.** Beta-high gamma PBC at individual channels in LMFG, RMTG, and LMTG. Crimson lines indicate significant differences from the mean of 400-600 ms of fixation (black rectangle at the bottom; $p < 0.05$, cluster test; 200 ms running window, 50% overlap). A 200 ms smoothing window was used for visualization but not for testing. Shaded regions represent ±1 SEM.

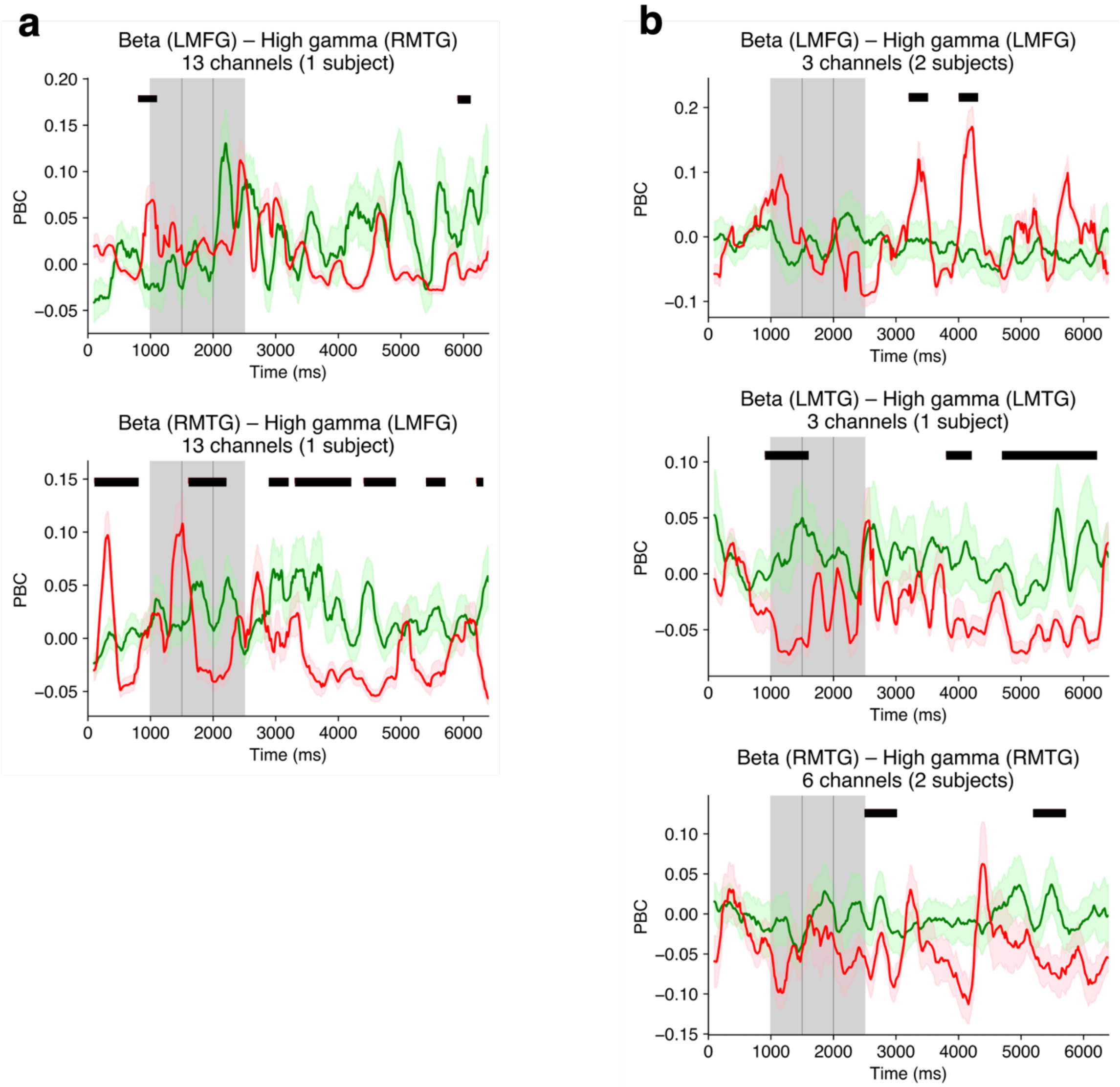


**Figure S11. Channels with elevated delay-period PBC differentiates performance | a.** Trial-balanced correct versus incorrect beta- high gamma PBC for channels with highest delay-period beta-high gamma PBC of LMFG-RMTG during correct trials (top 20%). **b.** Same as **a** for channels within LMFG, LMTG and RMTG. Black lines indicate significant differences between correct and incorrect trials ($p < 0.05$, cluster test; 200 ms running window, 50% overlap). A 200 ms smoothing window was used for visualization but not for testing. Shaded regions represent ±1 SEM.

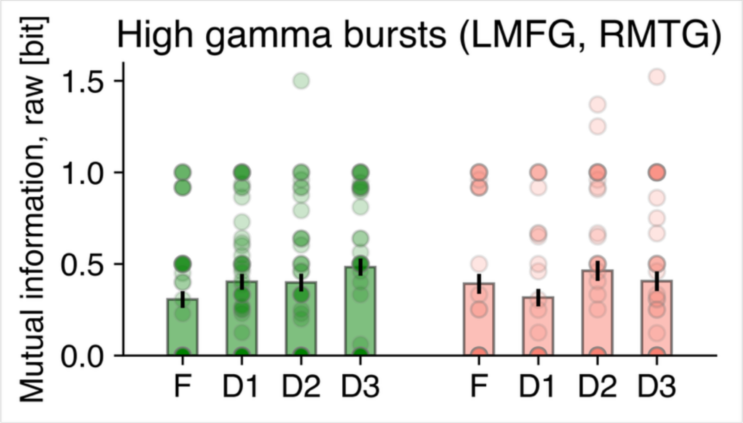


**Fig. S12. Mutual information between LMFG and RMTG high gamma bursts during trial, averaged over trials (66 channel pairs, data from 2 subjects) |** Bars show per-trial mutual information without baseline-subtraction computed over 100-900 ms of fixation (F) and three sub-periods of the delay period (D1: 2600-3400 ms, D2: 4000-4800 ms, D3: 5200-6000 ms). No significant differences found (2-way GLME, post hoc pairwise tests, Tukey correction).

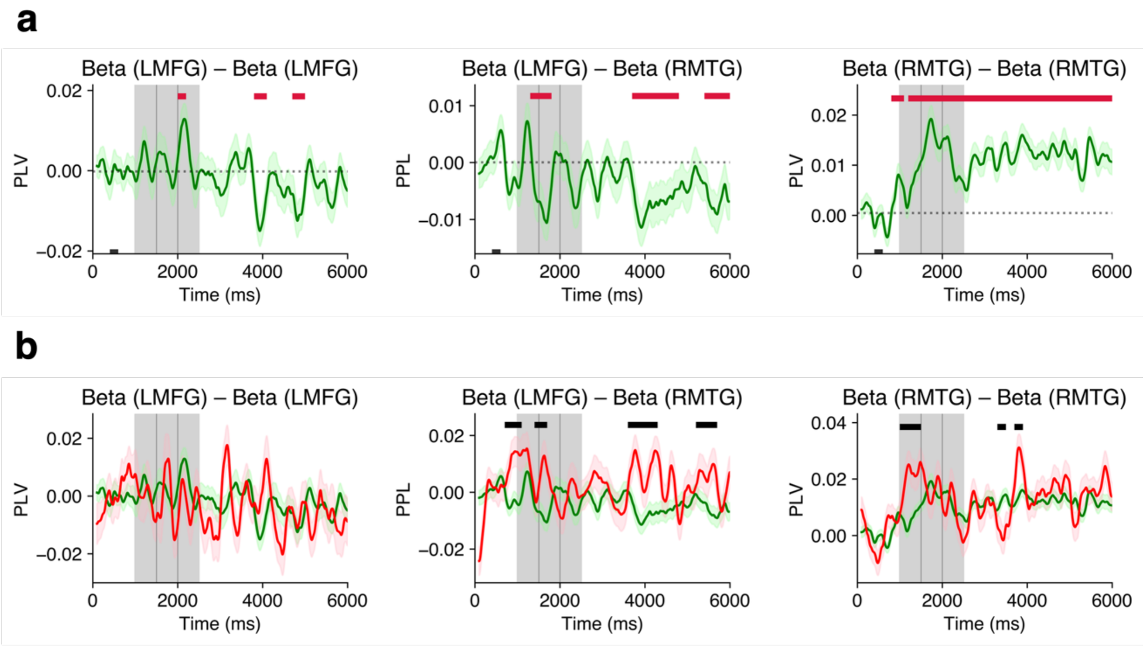


**Fig. S13. Beta-band phase-phase locking (PPL) in LMFG and RMTG (42 channel pairs within LMFG, data from 3 subjects; 66 pairs between LMFG and RMTG, data from 2 subjects; 166 pairs within RMTG, data from 5 subjects) | a.** Baseline-subtracted beta-band PLV and PPL for correct trials, mean over channels. Frontotemporal beta PPL decreased during the delay period relative to fixation. Beta PLV within the temporal region increased during the delay period. **b.** Baseline-subtracted beta-band PLV and PPL for correct compared to incorrect trials, mean over channels. In panel **a**, crimson lines indicate significant differences from the mean of 400-600 ms of fixation indicated by black rectangle at the bottom ($p < 0.05$, cluster test; 200 ms running window, 50% overlap). In panel **b**, black lines at the top indicate significant differences between correct and incorrect trials ($p < 0.05$, cluster test; 200 ms running window, 50% overlap). In panels **a** and **b**, the baseline period was 100-900 ms from the start of the trial. A 200 ms smoothing window was used for visualization but not for testing. Shaded regions represent ±1 SEM.

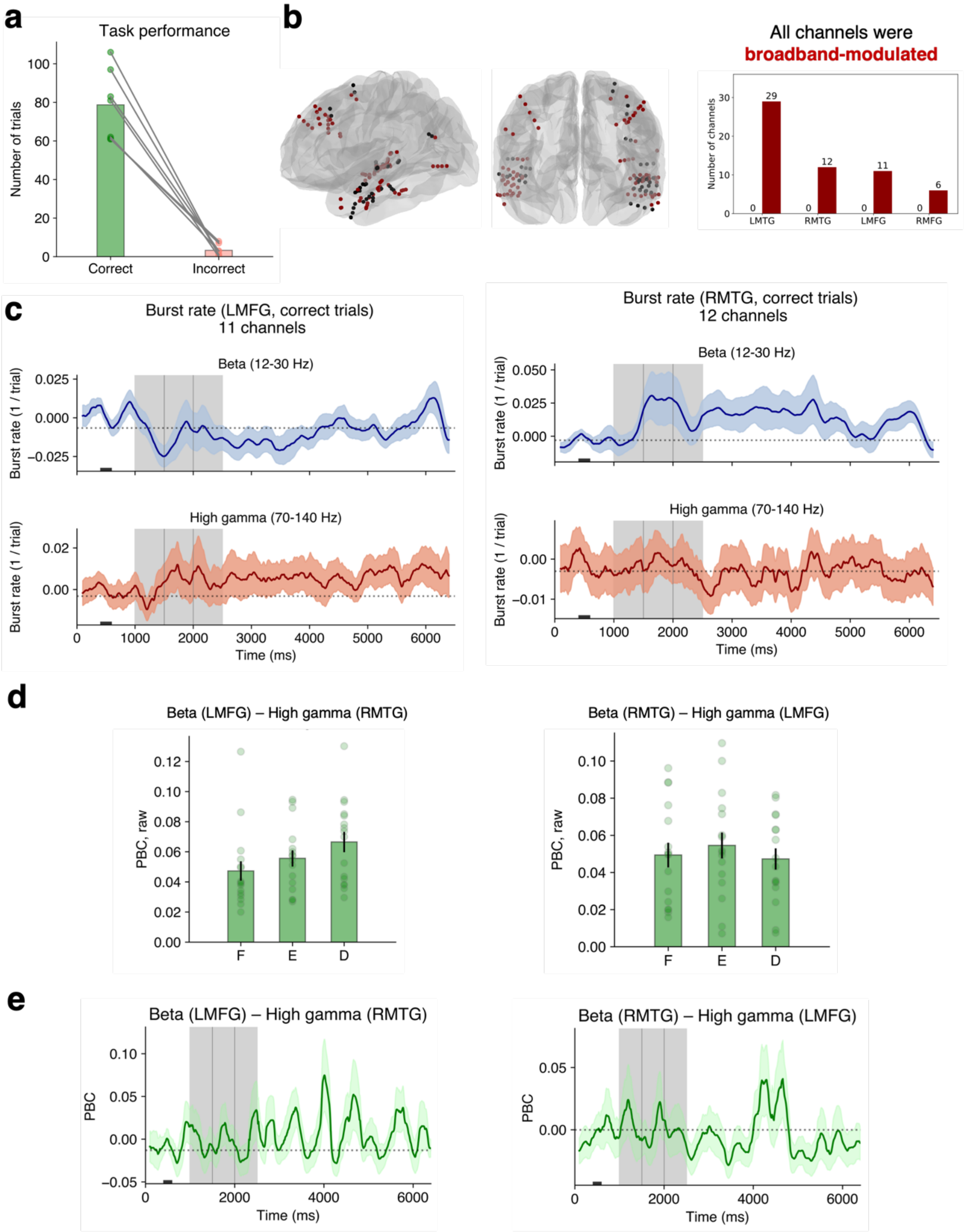


**Fig. S14. Results from the University of Missouri cohort, part 1 | a.** Task performance across 6 subjects. The number of incorrect trials was not sufficient to perform performance comparisons. **b.** Electrode locations across patients and the number of broadband-modulated channels across ROIs. **c.** Burst rates in LMFG and RMTG (correct trials). **d.** Data from 3 subjects, raw period-averaged PBC in correct trials (16 channel pairs between LMFG and RMTG). Error bars correspond to ±1 SEM. **e.** Data from 3 subjects, temporal dynamics of the baseline-subtracted PBC in correct trials. In panels **c** and **e**, crimson lines indicate significant differences from the mean of 400-600 ms of fixation (black rectangle at the bottom; $p < 0.05$, cluster test; 200 ms running window, 50% overlap). A 200 ms smoothing window was used for visualization but not for testing. Shaded regions represent ±1 SEM.

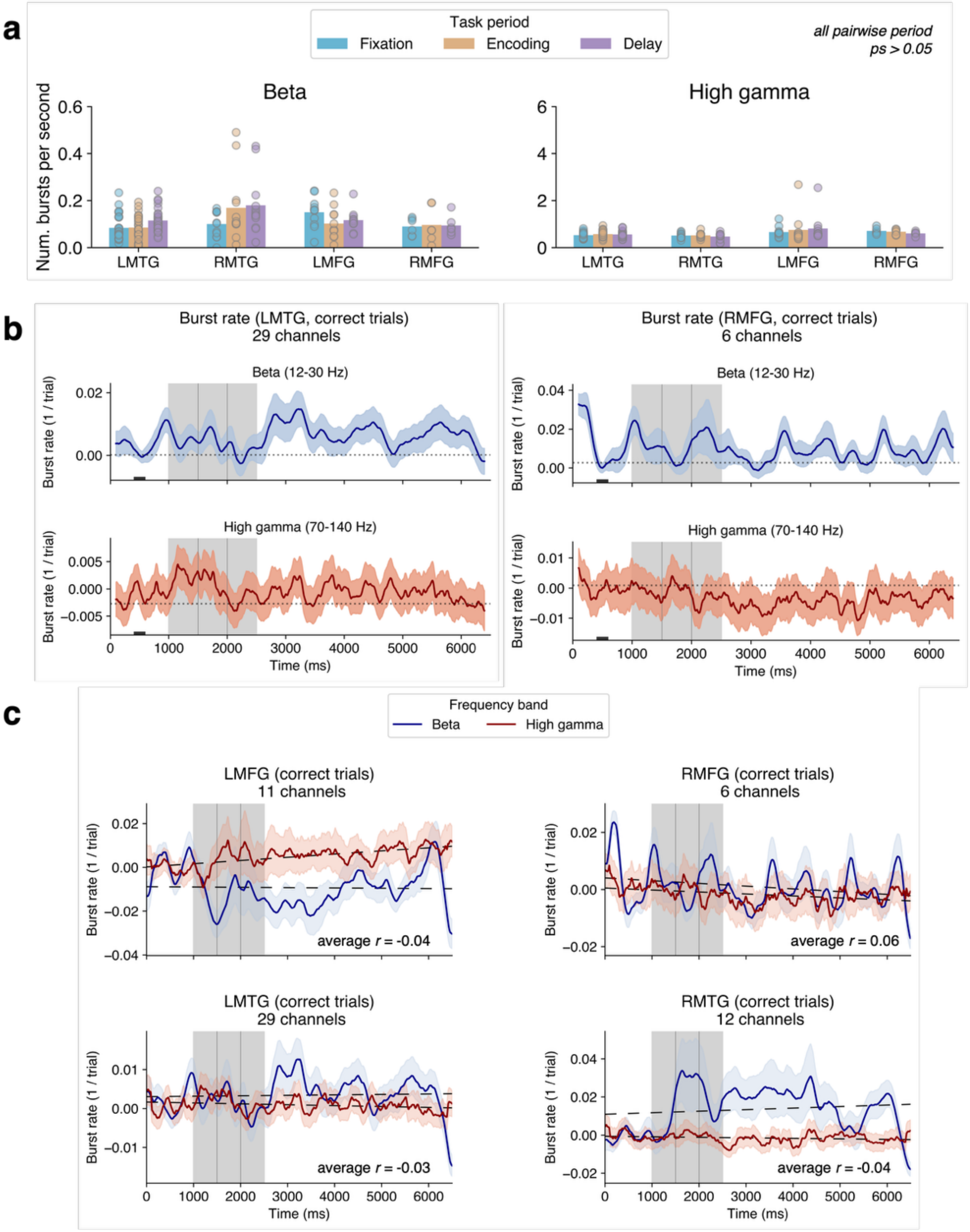


**Fig. S15. Results from the University of Missouri cohort, part 2 | a.** Number of bursts per second across periods and ROIs. No significant differences between periods were found in any ROI (post hoc pairwise tests, Tukey correction). **b.** Burst rates in LMFG and RMTG (correct trials). Crimson lines indicate significant differences from the mean of 400-600 ms of fixation (black rectangle at the bottom; $p < 0.05$, cluster test; 200 ms running window, 50% overlap). **c.** Correlation between high gamma and beta burst rates across ROIs. As in the Utah cohort, significant temporal anti-correlation was found in LMFG (in 9/11 channels), LMTG (in 23/29 channels), RMTG (in 8/12 channels). Conversely, RMFG showed significant positive correlation (in 6/6 channels). In panels **b** and **c**, a 200 ms smoothing window was used for visualization but not for testing. Shaded regions represent ±1 SEM.

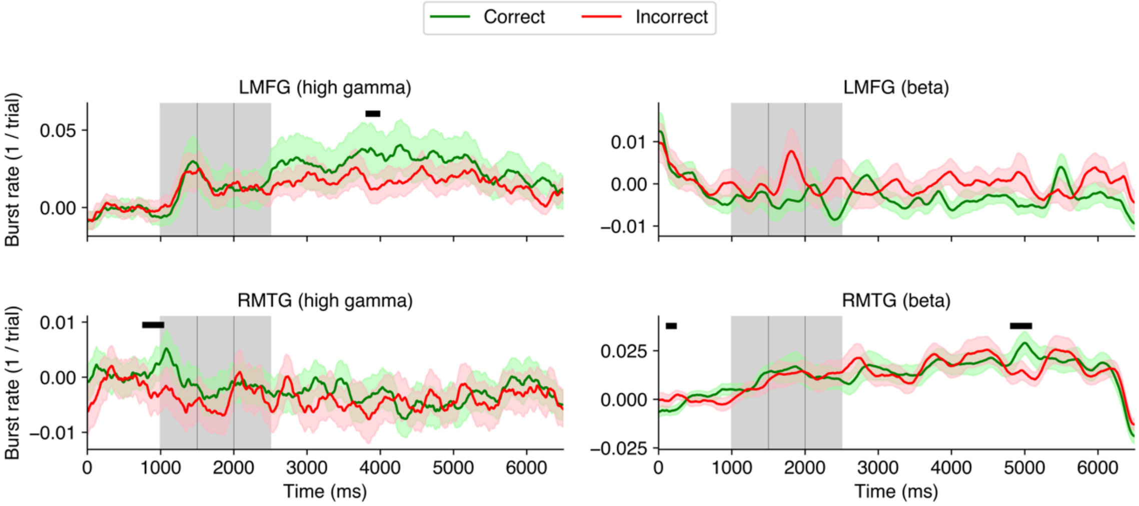


**Fig. S16. Burst rates in LMFG and RMTG scale with number correct |** Here, ‘incorrect’ trials were defined as those with 0, 1, or 2 (i.e., <3) items chosen correctly. Thus, this is close in performance to the ‘correct’ trials, in which all 3 items were correct. Mean LMFG high gamma and RMTG beta burst rates for ‘<3 items correct’ condition was more similar ‘3 correct’ condition versus the ‘<2 items correct’ condition shown in **Figure 4** of main text. This implies that burst rates scale with memory performance. Shaded regions represent ±1 SEM. The black bars correspond to periods of significant difference between correct and incorrect averages over channels ($p < 0.05$, cluster test; 200 ms running window, 75% overlap). The baseline period used is 200-800 ms from the start of the trial. A 200 ms smoothing window was used for visualization but not for testing.

**Table S1. Patient demographics and relevant characteristics (University of Missouri cohort)**

| Patient | Age | Gender | Handedness | Language laterality | Epilepsy duration (years) | ROI of SOZ | Number of channels contributed to the analyses |
|---|---|---|---|---|---|---|---|
| MU1 | 25 | Female | Right | Left | 3 | Left hippocampus | 6 |
| MU2 | 41 | Female | Right | | 13 | Left lingual gyrus and fusiform gyrus | 9 |
| MU3 | 31 | Female | Right | Left | 1 | Right hippocampus | 5 |
| MU4 | 46 | Female | Right | Left | 25 | Right hippocampus, Left middle temporal gyrus, Left hippocampus | 7 |
| MU 5 | 50 | Female | Right | Left | 23 | Left hippocampus and left fusiform gyrus | 11 |
| MU 6 | 29 | Male | Right | Left | 7 | Right frontotemporal area | 12 |

**Table S2. Laterality of patients' channels (University of Missouri cohort)**

| Patient | LMFG | RMFG | LMTG | RMTG |
|---|---|---|---|---|
| MU1 | 1 | | 4 | 1 |
| MU2 | | | 8 (s1) / 9 (s2) | |
| MU3 | | 2 | 3 | |
| MU4 | 5 | | | 2 |
| MU5 | 5 | | 5 | 1 |
| MU6 | | 4 | | 8 |

s: session

**References**


1. E. D. Gribkova, B. A. Ibrahim, D. A. Llano, A novel mutual information estimator to measure spike train correlations in a model thalamocortical network. *J Neurophysiol* **120**, 2730–2744 (2018)
2. E. Rezayat, *et al.*, Frontotemporal coordination predicts working memory performance and its local neural signatures. *Nat Commun* **12**, 1103 (2021).
